# Understanding the interaction of 14-3-3 proteins with *h*DMX and *h*DM2: a structural and biophysical study

**DOI:** 10.1101/2021.12.17.473238

**Authors:** Sonja Srdanovic, Madita Wolter, Chi H. Trinh, Christian Ottmann, Stuart L. Warriner, Andrew J. Wilson

## Abstract

p53 plays a critical role in regulating diverse biological processes: DNA repair, cell cycle arrest, apoptosis, and senescence. The p53 pathway has therefore served as the focus for drug-discovery efforts. p53 is negatively regulated by *h*DMX and *h*DM2; prior studies have identified 14-3-3 proteins as *h*DMX and *h*DM2 client proteins. 14-3-3 proteins are adaptor proteins that modulate localisation, degradation and interactions of their targets in response to phosphorylation. Thus 14-3-3 proteins may indirectly modulate the interaction between *h*DMX or *h*DM2 and p53 and represent potential targets for modulation of the p53 pathway. In this manuscript we report on the biophysical and structural characterization of peptide/protein interactions that are representative of the interaction between 14-3-3 and *h*DMX or *h*DM2. The data establish that proximal phosphosites spaced ∼20-25 residues apart in both *h*DMX and *h*DM2 co-operate to facilitate high-affinity 14-3-3 binding and provide structural insight that can be utilized in future stabilizer/inhibitor discovery efforts.

## Introduction

The transcription factor p53 is often described as “the guardian of the genome” because of its central role in regulating diverse biological processes: DNA repair, cell cycle arrest, apoptosis, and senescence.^1^ Thus, the p53 pathway has seen significant attention in developing new anticancer treatments.^2^ One such target has been the interaction between p53 and *h*DM2 or *h*DMX (Fig. 1a).^3-5^ p53 is negatively regulated by *h*DMX and *h*DM2;^6-10^ they bind to the transactivation domain of p53 sterically blocking its interaction with DNA,^9-12^ p53 is additionally targeted for nuclear export and proteasomal degradation through *h*DM2 which bears an E3 ligase domain.^13-15^ Although *h*DMX lacks such a domain, the E3 ligase activity of *h*DM2 is enhanced through formation of *h*DMX/*h*DM2 heterodimers.^16-18^ This continuous targeting of p53 for degradation keeps p53 levels low under normal conditions. It is only under stressed conditions (e.g. DNA damage, hypoxia, oncogene activation and ribosomal stress) that p53 is rapidly activated through multiple post translational modifications; subsequent stabilization and accumulation of p53 upregulates gene expression leading to cell cycle arrest and/or apoptosis. This tightly controlled autoregulatory loop is thus of vital importance; knock out studies of the oncogenes MDMX and MDM2 (the murine homologues of *h*DMX and *h*DM2) have shown that these proteins are essential in mice studies.^19^ However, *h*DM2 and *h*DMX are themselves subject to regulatory control and have additional independent roles in cell cycle regulation.^20-22^ For instance, in addition to modulating p53 binding, post translational modification of *h*DMX and *h*DM2 such as phosphorylation influences their localization and degradation.^21-25^

**Figure 1.**
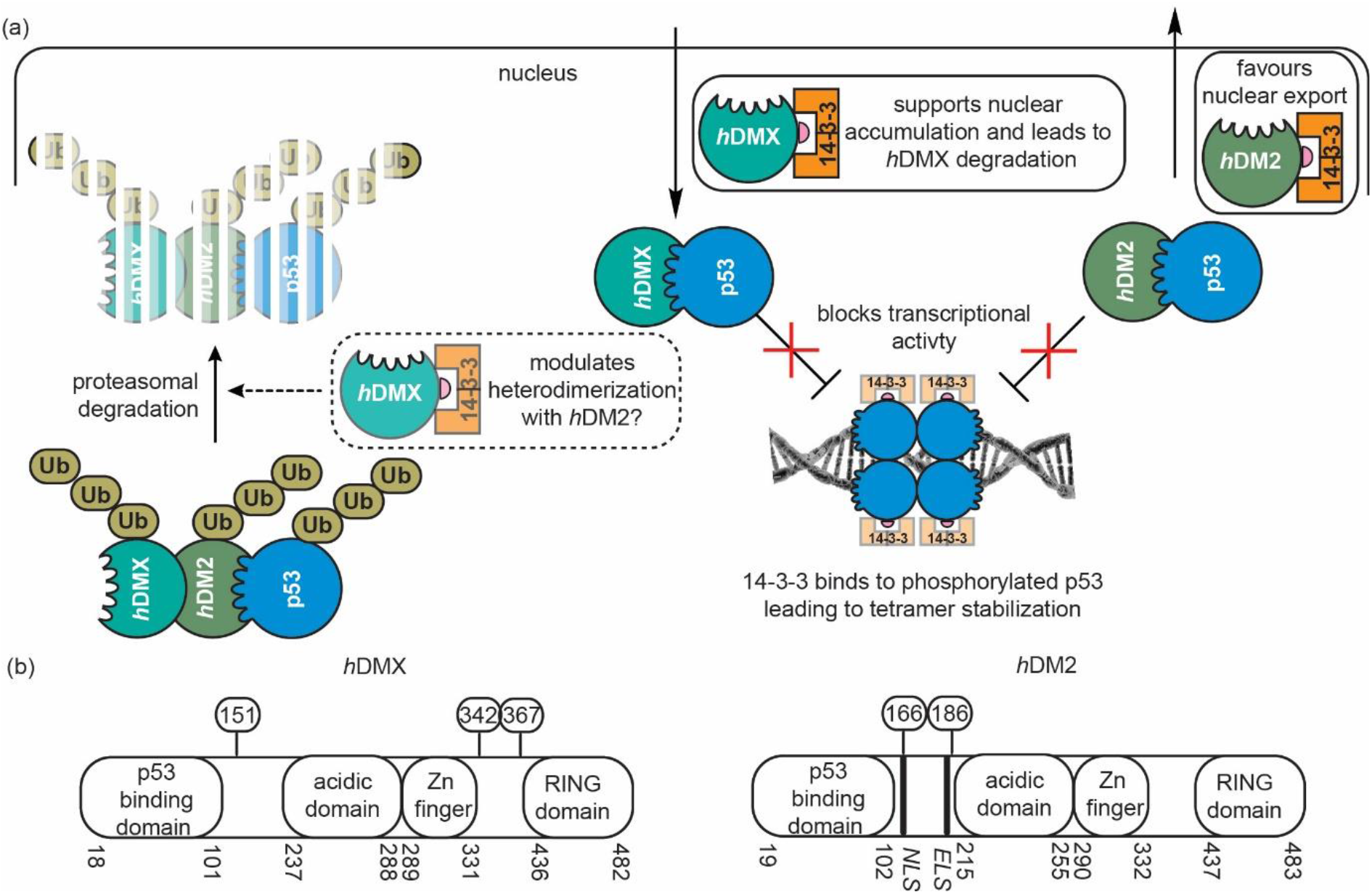
Negative regulation of the p53 pathway by *h*DMX and *h*DM2: (a) *h*DMX (turquoise) and *h*DM2 (green) bind p53 (blue) and subsequently inhibit its transcriptional activity. *h*DM2 also facilitates nuclear export of p53, and as a ubiquitin ligase additionally tags p53 for degradation, whilst *h*DMX accelerates *h*DM2 mediated ubiquitin ligase activity. 14-3-3 (orange) interacts with: (i) phosphorylated p53 to stabilize tetramer formation, (ii) phosphorylated *h*DMX leading to its nuclear accumulation and degradation, and, possibly modulation of its interaction with *h*DM2, (iii) *h*DM2 to facilitate its nuclear export. (b) domain structure of *h*DM2 and *h*DMX highlighting phosphosites on *h*DM2 and *h*DMX investigated in this work: Ser342 and Ser367 in intracellular localization, protein degradation and ubiquitination, and additionally Ser367 is associated with cell growth and protein stabilization. Ser166 and Ser186 on *h*DM2 are involved in apoptosis, signalling pathway regulation and ubiquitination.

Several kinases have been reported to phosphorylate *h*DMX and/or *h*DM2 whilst cellular studies have shown that phosphorylated *h*DMX and *h*DM2 interact with 14-3-3.^26^ First identified in 1967,^27^ 14-3-3 proteins are eukaryotic adapter proteins that control diverse physiological functions through interaction with a large network of proteins e.g. they are involved in signal transduction, protein trafficking, apoptosis, and cell cycle regulation.^28^ Given these functions, they have been shown to play an important role in inflammatory conditions, neurodegenerative diseases, cancer and cystic fibrosis.^29-33^ and have consequently received attention as potential drug-discovery targets.^28, 34^ The 14-3-3 family of proteins consist of seven isoforms in humans: β, γ, ε, η, ζ, σ and τ/θ, all of which share a high degree of sequence conservation (see supporting information, Fig. S1). Even though sequence conservation between the isoforms can indicate redundancy in protein function, this is not the case for 14-3-3 proteins as different affinities towards a target protein are clear between the isoforms.^35-36^ 14-3-3 proteins exist as homo or heterodimers where each monomer consists of nine α-helices forming an amphipathic binding groove. Four α-helices are directly responsible for dimer formation, while the rest of the α-helices form the “W” shape of the protein dimer.^37^ The N terminal domain of 14-3-3 proteins modulates homo/heterodimerization while the C terminal domain binds client proteins.^37^ Activity and modulation of many cellular pathways by 14-3-3 is controlled by phosphorylation events. Target proteins are typically recognized by 14-3-3 through well defined sequences that can be generalized into three categories: I.) RXX pS/pT XP, II.) RX(Y/F)X pS/pT XP, III.) XX pS/pT X COOH (pS and pT being phosphorylated Ser and Thr, respectively, X being any amino acid).^38^ In mode I and II, proline is always located at position +2 in relation to the central pSer/pThr and arginine is at position -3 in mode I. Mode III represents C terminal sequences where pSer/pThr are the penultimate amino acid of the partner protein.

Phosphorylated residues on *h*DMX or *h*DM2 considered relevant to recognition by 14-3-3 proteins, as well as their role in cell signalling are outlined in Fig. 1b.^26, 39-43^ Relevant sites on *h*DMX: Ser342 and Ser367 are targeted by several kinases (e.g. AMP,^43^ Akt,^39^ Chk2^26, 42^ or Chk1^40^). 14-3-3 phosphorylation in response to stress or DNA damage with consequent enhancement of *h*DM2 mediated *h*DMX degradation has been reported,^42, 44^ together with a role for Ser367.^41^ Phosphorylation of *h*DMX on Ser367 by Chk2 was shown to promote 14-3-3 binding, *h*DMX nuclear import and degradation by *h*DM2.^26^ Similarly, Chk1 was shown to phosphorylate *h*DMX at Ser367, to promote 14-3-3γ/hDMX interaction and cytoplasmic retention of *h*DMX to counter *h*DMX-enhanced p53 ubiquitination, leading to its stabilization and activation.^40^ However, the potential relevance of both Ser342 and Ser367 to interaction with 14-3-3 has been noted.^42, 45^ In contrast, Akt has been shown to phosphorylate *h*DMX on Ser367, promoting 14-3-3 binding, stabilizing *h*DMX and consequently enhancing *h*DM2 stability leading to suppression of p53 transcriptional activity.^39^Adenosine monophosphate (AMP) mediated phosphorylation of *h*DMX on Ser342 has been shown to induce *h*DMX/14-3-3 interaction resulting in inhibition of p53 ubiquitylation and consequent stabilization.^43^ *h*DM2 phosphorylation in the NLS/ELS region can lead to its stabilization by PKB/Akt,^46-47^ although the effects are complex; phosphorylation on Ser166 and Ser188 are promoted by Akt kinase,^48^ and result in positive regulation of *h*DM2 by inhibiting its self-ubiquitination and nuclear translocation.^46, 49-50^ Phosphorylation at Ser166 and Ser188 does not appear to result in binding of *h*DM2 to 14-3-3^51^ whilst such an interaction was observed when phosphorylated at Ser166 and Ser186 by Pim kinase, which simultaneously suppresses phosphorylation of Ser188.^51^ Finally, interaction between *h*DM2 phosphorylated in the ring domain and 14-3-3σ was observed to accelerate the self ubiquitination of *h*DM2, with a resulting downstream stabilization of p53.^52^ Thus cellular studies have linked *h*DMX and *h*DM2 to 14-3-3 proteins; 14-3-3 proteins may indirectly modulate the interaction between *h*DMX or *h*DM2 and p53 and represent potential targets for pharmacological modulation of p53^21, 23^ but the molecular bases of these interactions remain unexplored. In this manuscript we report on the biophysical and structural characterization of peptide/protein interactions that are representative of the interaction between 14-3-3 and *h*DMX or *h*DM2. The data reveal the interactions to be mode I;^53-54^ whilst such 14-3-3 interactions are not stabilized by typical 14-3-3 interaction stabilizers e.g. Fusicoccin and Cotelynin A.^55^ These data provide structural insights that could be utilized in future stabilizer/inhibitor discovery efforts. More significantly, our results reveal that in both cases proximal phosphosites spaced ∼20-25 residues apart in both *h*DMX and *h*DM2 co-operate to facilitate high-affinity 14-3-3 binding and do so through 1:1 stoichiometry.

## Results

There are multiple serine and threonine residues in the *h*DMX and *h*DM2 proteins that could be phosphorylated (protein sequences with highlighted Ser and Thr residues on *h*DMX and *h*DM2 proteins, as well as their alignment can be found in the supporting information Fig. S2-3). Therefore, *14-3-3Pred*^56^ was used to identify sequences from *h*DM2 and *h*DMX that might serve as 14-3-3 binding sequences. *14-3-3Pred* is a webserver that predicts 14-3-3 binding sites in client proteins based on published examples of 14-3-3 interactions with specific amino acids surrounding Ser/Thr. Scores are assigned to each Ser/Thr residue on the protein target whose sequences in FASTA format are queried, using an ensemble approach comprising three different base classification systems: position specific scoring matrix (PSSM), support vector machines (SVM) and artificial neural network (ANN). End score values are calculated as the average of three independent methods with a score >0.5 indicating high probability of the sequence binding to 14-3-3. The webserver also provides information on the phosphorylation state for each Ser/Thr residue. The phosphorylated sites on *h*DMX and *h*DM2 shown to be relevant for interaction of 14-3-3 in previous cell-based studies^26, 40^-^43, 48, 51^ were predicted to have a high probability of 14-3-3 binding, except pSer342 on *h*DMX, which showed low probability of binding (score 0.19) whereas Ser151 on *h*DMX – a previously unreported site of phosphorylation - was predicted to be a potential 14-3-3 binding site (score 0.75). Using the *14-3-3Pred* results server several key regions of *h*DMX and *h*DM2 were thus identified. Peptides containing these binding regions were designed bearing five/six residues flanking the central phosphorylated residue on either side and prepared by solid-phase peptide synthesis (SPPS). Given the proximity of pSer342 and pSer367 from *h*DMX, as well as pSer166 and pSer186 from *h*DM2, we also prepared doubly phosphorylated sequences for both to probe for avidity effects. Peptides were synthesized using a combination of manual or automated Fmoc SPPS on rink amide resin (Rink amide ProTide for doubly phosphorylated sequences) and modified at the N terminus by acetylation for isothermal titration calorimetry (ITC), surface plasmon resonance (SPR) and crystallography studies or with 5,6-carboxyfluorescein following an Ahx linker to function as a tracer in fluorescence anisotropy (FA) assays (see ESI for further details).

### Biophysical characterization of the interaction of *h*DMX and *h*DM2 peptides with 14-3-3

To determine the binding affinity between all 7 isoforms of 14-3-3 proteins and *h*DMX and *h*DM2 peptides, FA experiments were carried out. 14-3-3 proteins were titrated against a constant concentration of the tracer resulting in an increase of anisotropy upon engaging 14-3-3 proteins (Fig. 2a-c for 14-3-3 binding *h*DMX sequences and Fig. S4-6, for 14-3-3 binding *h*DM2 sequences, non-binding sequence and unphosphorylated controls). A noteworthy feature of the data is that different maximum anisotropy levels were observed between *h*DMX or *h*DM2 peptides and 14-3-3 isoforms, indicating different diffusion properties of the fluorophore when bound to the different 14-3-3 isoforms. This may: (i) reflect differences in the interaction between fluorophore and protein restricting the former’s mobility to a greater or lesser degree between isoforms, or (ii) arise as a consequence of a more complex equilibrium e.g. isoform dependent variation in 14-3-3 dimerization^57^ across the concentration range of the assay. This will be investigated in future studies. The consequence of this behaviour is that for weaker binding peptides where saturation was not observed, it was not possible to determine a *K*_d_ value. Thus, in some cases, e.g. *h*DM2_180-192_^pSer186^, K_d_ values (see below) with τ, σ and ε could only be estimated to be greater than limiting values. Similarly, accurate fitting for doubly phosphorylated peptides could be obtained only for η, γ and β isoforms. Peptides *h*DMX_335-349_^pSer342^ and *h*DMX_361-374_^pSer367^ showed affinity for 14-3-3 proteins, whereas peptide *h*DMX_144-158_^pThr151^ exhibited no response under the conditions of the assay (see supporting information Fig S5). None of the corresponding unphosphorylated control *h*DMX sequences exhibited evidence of binding under the conditions of the assay confirming the role of phosphorylation for binding of pSer342 and pSer367 to 14-3-3 (see supporting information Fig. S6). *h*DM2 peptides representing sites pSer166 and pSer186 showed low micromolar affinities to all 14-3-3 isoforms (supporting information, Fig S4).

**Figure 2.**
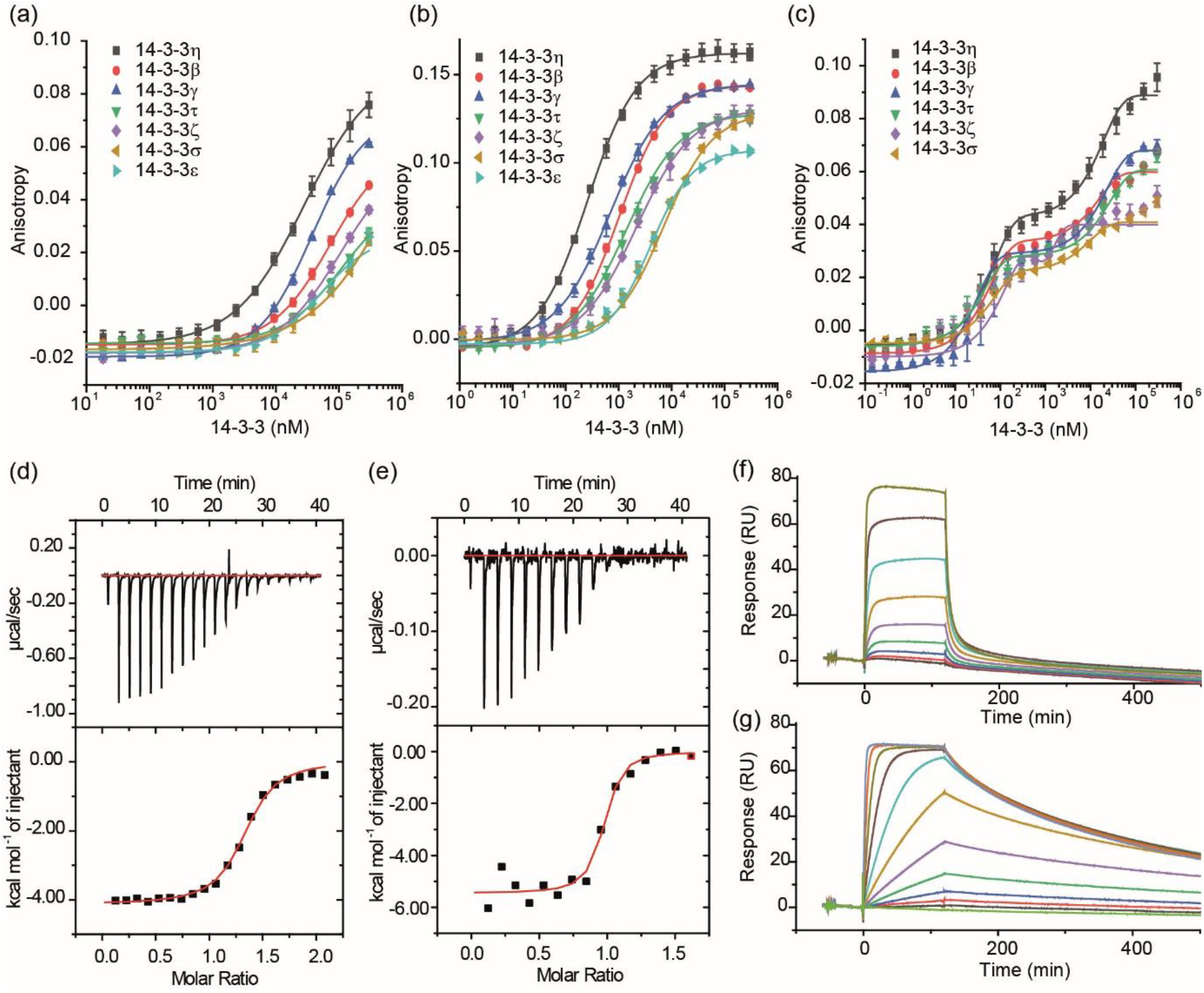
Biophysical analyses of the interaction between 14-3-3 and *h*DMX peptides (a-c) FA assays for the *h*DMX peptides (a) *h*DMX_335-349_^pSer342^ (b) *h*DMX_361-374_^pSer367^ and (c) *h*DMX _335-373_^pSer342/pSer367^ with all isoforms of 14-3-3 (FAM tracer peptide 50 nM, in 10 mM HEPES, 150 mM NaCl, 0.1% Tween 20, 0.1% BSA, pH 7.4, concentration of proteins given as 14-3-3 monomer concentration); (d-e) ITC data for the interaction of *h*DMX peptides with 14-3-3η; (d) *h*DMX_361-374_^pSer367^; (e) *h*DMX_335-373_^pSer342/pSer367^ (peptides were titrated into 14-3-3η {0.1M for mono phosphorylated peptide and 0.02M for doubly phosphorylated peptide}, 25°C, 25 mM HEPES pH 7.5, 100 mM NaCl, 10 mM MgCl_2_, 0.5 mM TCEP); (f-g) Dose response SPR experiments for the binding of *h*DMX peptides to immobilized 14-3-3η (f) *h*DMX_361-374_^pSer367^; (g) *h*DMX _335-373_^pSer342/pSer367^ (peptides – concentration at 10x the K_d_ for each peptide – were passed over immobilized 14-3-3η, 25°C, 25 mM HEPES pH 7.5, 100 mM NaCl, 10 mM MgCl_2_; experiments were performed in a multicycle kinetic format and data was fitted to a Langmuir model. K_d_ values were determined by fitting maximal response level at the end of injection against protein concentration using a steady state affinity model in the Biocore evaluation software.

**Figure 3.**
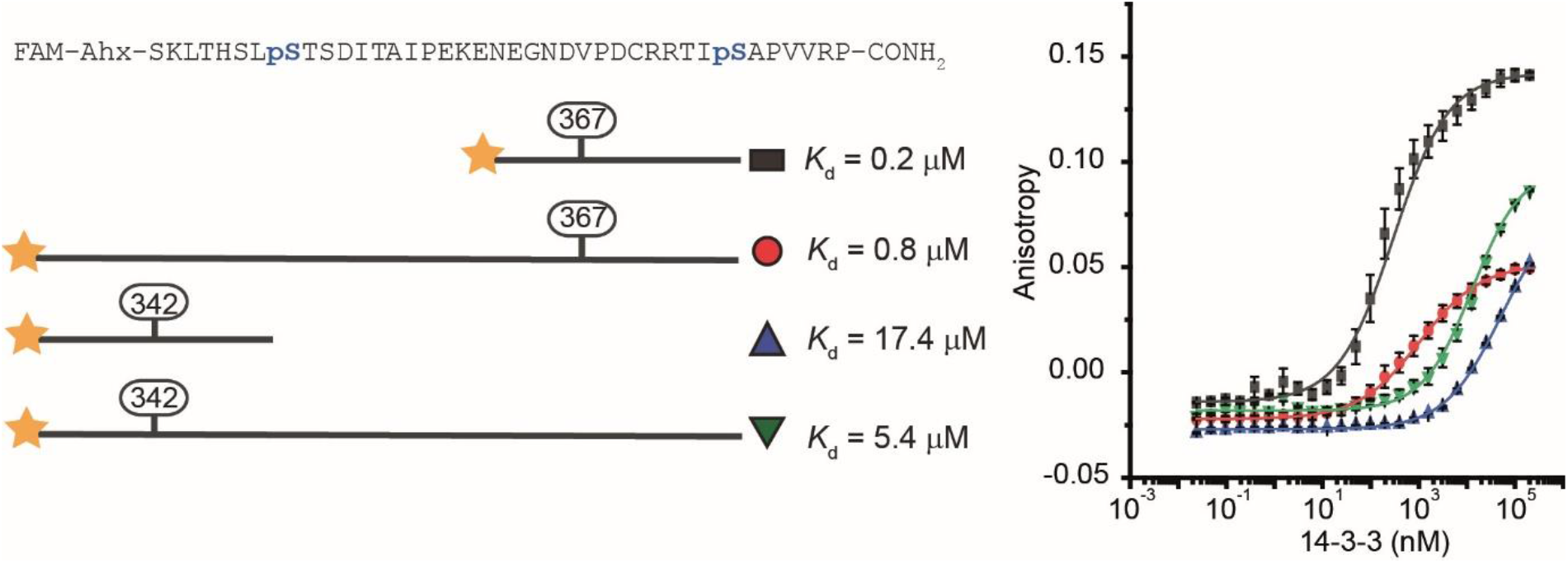
Difference in the affinity between short *h*DMX_335-349_^pSer342^ and *h*DMX_361-374_^pSer367^ peptides and their longer singly phosphorylated variants (*h*DMX_335-373_^pSer342^ and *h*DMX _335-373_^pSer367^) as determined by fluorescence anisotropy titration (tracer 50 nM, 10 mM HEPES, 150 mM NaCl, 0.1% Tween 20)

Generally, all peptides exhibited tightest binding towards the 14-3-3η isoform, followed by γ, β, τ, ζ, σ and ε, respectively. *h*DMX and *h*DM2 peptides mimicking one phosphorylation site fit well to a 1:1 binding isotherm. In this previously described model each 14-3-3 dimer protein can bind two peptides, one peptide per 14-3-3 protomer.^37^ *K*_d_ values for all 14-3-3 isoforms are reported in Table 2, but for simplicity only the 14-3-3η isoform is discussed below. *h*DMX_361-374_^pSer367^ exhibited higher affinity (*K* _d_ = 98.8 ± 4.6 nM) than *h*DMX_335-349_^pSer342^ (*K*_d_ = 20.0 ± 1.2 μM). *h*DM2 sequences bound with weaker affinity (*h*DM2_160-171_^pSer166^ *K*_d_ = 10.2 ± 0.3 μM and *h*DM2_180-192_^pSer186^ *K*_d_ = 31.6 ± 1.9 μM). As might be expected the highest affinity sequence (*h*DMX_361-374_^pSer367^) is a mode I 14-3-3 sequence/ binder (see later) with Pro at the +2 position and Arg at the -3 position.

**Table 1.**
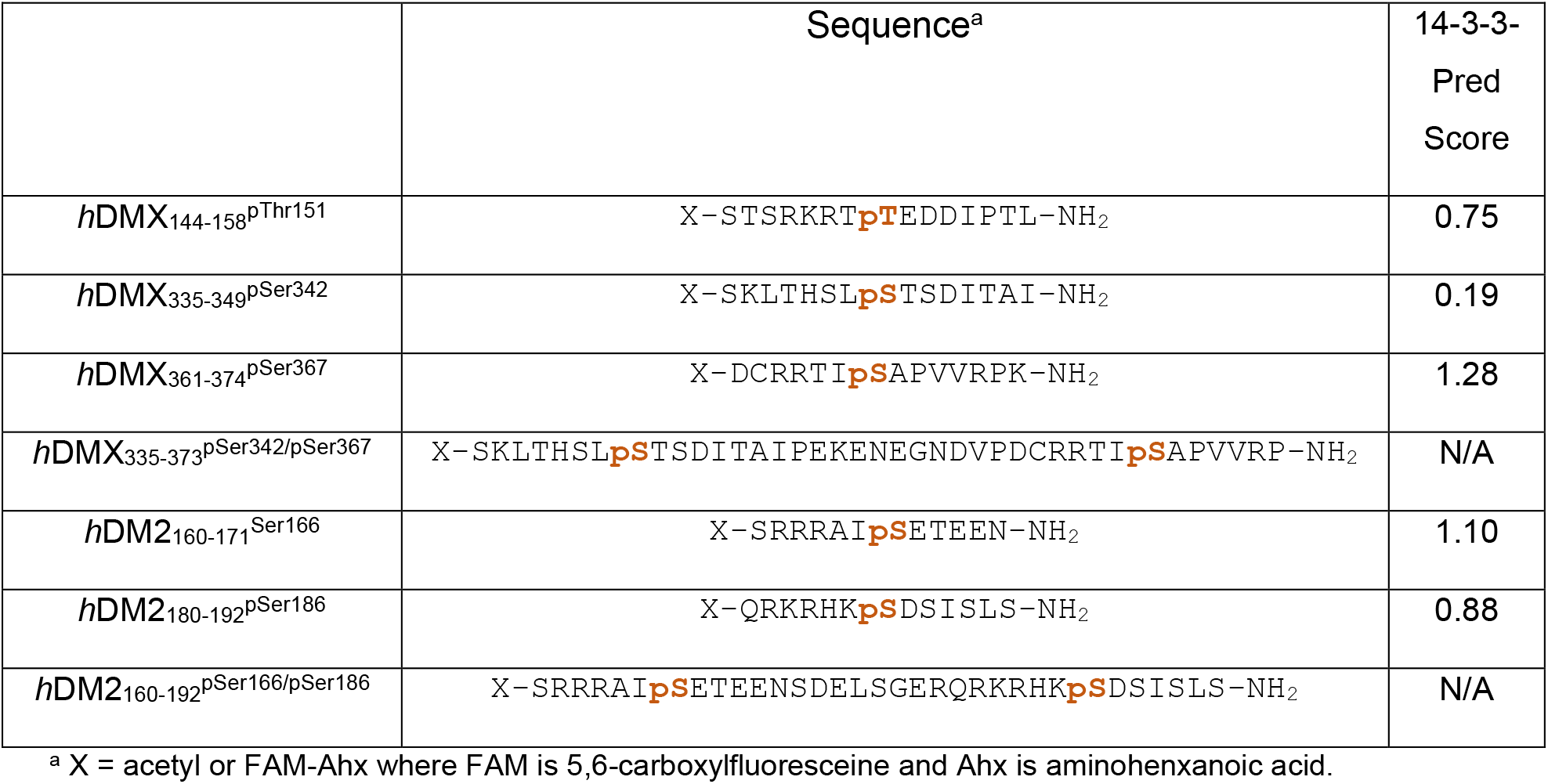
14-3-3 binding *h*DM2 and *h*DMX Sequences studied

|  | Sequence<sup>a</sup> | 14-3-3-Pred Score |
| --- | --- | --- |
| *hDMX*<sub>144-158</sub><sup>pThr151</sup> | X-STSRKRT**pT**EDDIPTL-NH<sub>2</sub> | 0.75 |
| *hDMX*<sub>335-349</sub><sup>pSer342</sup> | X-SKLTHSL**pS**TSDITAI-NH<sub>2</sub> | 0.19 |
| *hDMX*<sub>361-374</sub><sup>pSer367</sup> | X-DCRRTI**pS**APVVRPK-NH<sub>2</sub> | 1.28 |
| *hDMX*<sub>335-373</sub><sup>pSer342/pSer367</sup> | X-SKLTHSL**pS**TSDITAIPEKENEGNDVPDCRRTI**pS**APVVRP-NH<sub>2</sub> | N/A |
| *hDM2*<sub>160-171</sub><sup>Ser166</sup> | X-SRRRAI**pS**ETEEN-NH<sub>2</sub> | 1.10 |
| *hDM2*<sub>180-192</sub><sup>pSer186</sup> | X-QRKRHK**pS**DSISLS-NH<sub>2</sub> | 0.88 |
| *hDM2*<sub>160-192</sub><sup>pSer166/pSer186</sup> | X-SRRRAI**pS**ETEENSDELSEGERQKRHK**pS**DSISLS-NH<sub>2</sub> | N/A |

<sup>a</sup> X = acetyl or FAM-Ahx where FAM is 5,6-carboxylfluoresceine and Ahx is aminohexanoic acid.

**Table 2.** K_d_ values for *h*DMX and *h*DM2 peptides binding to 14-3-3 proteins (in μM) (increasing affinity is denoted by increasingly darker grey shading)

| protein peptide | 14-3-3 $\eta$ | 14-3-3 $\beta$ | 14-3-3 $\tau$ | 14-3-3 $\zeta$ | 14-3-3 $\gamma$ | 14-3-3 $\sigma$ | 14-3-3 $\epsilon$ |
| --- | --- | --- | --- | --- | --- | --- | --- |
| *hDMX*<sub>335-349</sub><sup>pSer342</sup> | 20.0 ± 1.2 | 49.4 ± 11.5 | 107.8 ± 6.0 | 95.8 ± 3.6 | 37.8 ± 0.7 | 135.9 ± 3.6 | 53.4 ± 2.9 |
| *hDMX*<sub>361-374</sub><sup>pSer367</sup> | 0.099 ± 0.005 | 0.51 ± 0.02 | 1.2 ± 0.2 | 0.92 ± 0.03 | 0.53 ± 0.04 | 3.6 ± 0.4 | 2.7 ± 0.2 |
| *hDMX*<sub>335-373</sub><sup>pSer342/pSer367</sup><br>$K_{d1}$ | 0.030 ± 0.006 | 0.026 ± 0.099 | - | - | 0.091 ± 0.015 | - | - |
| *hDMX*<sub>335-373</sub><sup>pSer342/pSer367</sup><br>$K_{d2}$ | 10.7 ± 1.2 | 7.6 ± 0.8 | - | - | 18.7 ± 0.7 | - | - |
| *hDM2*<sub>160-171</sub><sup>pSer166</sup> | 10.2 ± 0.3 | 11.1 ± 0.5 | 29.3 ± 2.4 | 19.6 ± 0.2 | 3.9 ± 0.3 | 32.7 ± 1.4 | 23.0 ± 1.2 |
| *hDM2*<sub>180-192</sub><sup>pSer186</sup> | 31.6 ± 1.9 | 207.9 ± 6.3 | > 250 | 165.8 ± 5.6 | 82.9 ± 2.1 | > 170 | > 260 |
| *hDM2*<sub>160-192</sub><sup>pSer166/pSer186</sup><br>$K_{d1}$ | 0.050 ± 0.012 | - | - | - | 0.089 ± 0.010 | - | - |
| *hDM2*<sub>160-192</sub><sup>pSer166/pSer186</sup><br>$K_{d2}$ | 17.5 ± 2.5 | - | - | - | 28.0 ± 0.8 | - | - |

Because of its dimer structure, 14-3-3 is known to bind multiple binding partners simultaneously, either by binding two different phosphorylated residues from the same target or possibly two phosphorylated sites within a multiprotein complex.^31, 58-60^ Thus, doubly phosphorylated *h*DMX and *h*DM2 peptides were used to asses the affects of proximal phosphorylation on affinity. Two state binding in FA was observed for both doubly phosphorylated *h*DMX and *h*DM2 peptides (Fig. 2c for *h*DMX peptide and Fig S4 for *h*DM2 peptide). Fitting for the observed biphasic-dose response curves was performed in the same manner as for the short peptides by fitting each binding event separately to a 1:1 model (one peptide to one 14-3-3 monomer) so as to obtain stepwise K_d_ values. Accurate fitting was possible for 14-3-3 η, γ and β isoforms as reported in Table 2 (for 14-3-3η, *h*DMX_335-373_^pSer342/pSer367^ *K*_d1_ = 30.3 ± 6.4 nM and *K*_d2_ = 10.67 ± 1.2 μM, *h*DM2_160-192_^pSer166/pSer186^; *K*_d1_ = 49.6 ± 12.1 nM and *K*_d2_ = 17.5 ± 2.5 μM). Approximately 200-fold higher affinity K_d1_ was observed for *h*DM2_160-192_^pSer166/pSer186^ compared to *h*DM2_160-171_^pSer166^ whilst a more modest 3-fold increase in binding affinity for *h*DMX_335-373_^pSer342/pSer367^ was observed in comparison to *h*DMX_361-374_^pSer367^ (although enhancement was more pronounced for binding to β and γ isoforms).

ITC was used as an orthogonal assay to confirm the binding affinities of *h*DMX and *h*DM2 peptides to 14-3-3η and to gain additional information about the stoichiometry of the interactions. Moreover, such experiments provide thermodynamic data and as a label free method require no immobilization/functionalization of any of the binding partners. Here data were obtained by titration of peptide into 14-3-3η protein (Fig 2d-e and supporting information Fig. S7). Thermodynamic parameters for interaction of 14-3-3 with *h*DMX and *h*DM2 peptides obtained by ITC are summarized in Table 3.

**Table 3.** Thermodynamic parameters for *h*DMX(2) peptides / 14-3-3 proteins

| | $K_d$<br>(ITC) | $\Delta G$<br>(kJ/mol) | $\Delta H$<br>(kJ/mol) | $-T\Delta S$<br>(kJ/mol) | n | $K_d$<br>(SPR) | $k_{on}$<br>(1/Ms) | $k_{off}$<br>1/s |
| --- | --- | --- | --- | --- | --- | --- | --- | --- |
| *hDMX*<sub>335-349</sub><sup>pSer342</sup> | 21 ± 3<br>μM | -27 | -21 ± 1 | -5 | 0.72 | 37<br>μM | - | - |
| *hDMX*<sub>361-374</sub><sup>pSer367</sup> | 870 ± 95<br>nM | -35 | -16 ± 0.2 | -18 | 1.28 | 189<br>nM | 3.89<br>$\times 10^5$ | 0.1208 |
| *hDMX*<sub>335-373</sub><sup>pSer342/pSer367</sup> | 14 ± 10<br>nM | -45 | -22 ± 0.7 | -23 | 0.94 | 8.6<br>nM | 1.51<br>$\times 10^6$ | 0.0048 |
| *hDM2*<sub>160-192</sub><sup>pSer166/pSer186</sup> | 151 ± 57<br>nM | -39 | -12 ± 0.6 | -27 | 0.94 | 155<br>nM | 1.45<br>$\times 10^6$ | 0.1215 |

ITC data showed similar trends for binding of *h*DMX peptides to 14-3-3η to those observed in the FA assays; binding affinities of *h*DMX_361-374_^pSer367^, *h*DMX_335-349_^pSer342^ and *h*DMX_335-373_^pSer342/pSer367^ were *K*_d_ = 870 ± 95 nM, *K*_d_ = 21 ± 3 μM and *K*_d_ = 14 ± 10 nM. An increase in binding affinity when both binding sites of *h*DM2 are present was also confirmed, with *K*_d_ = 151 ± 57 nM. The stoichiometry between doubly phosphorylated *h*DMX and *h*DM2 peptides and 14-3-3η was found to be approximately 0.94 indicating 1:1 binding model. If two phosphorylated sites from one peptide were binding simultaneously, the expected stoichiometry would be around 0.5. The contrasting absence of two binding events in the ITC data when compared to the FA experiments can be accounted for by the different experimental configurations. In the former, titration of peptide into protein means that protein begins in excess and becomes saturated with the highest affinity site, whereas for FA, peptide begins in excess and protein (and therefore monomer/dimer ratio) varies in concentration across the experiment Finally, association and dissociation rates for the binding of *h*DMX and *h*DM2 peptides to 14-3-3η were determined by SPR (Fig 2e-f and supporting information Fig. S8-9). Similar *K*_d_ values were confirmed from dose response SPR and where the affinities were calculated directly, from the k_on_/k_off_ values (for peptides where kinetics could be calculated). SPR measured affinities were broadly in agreement with *K*_d_ values determined from FA and ITC experiments (Table 2 and 3). For peptides exhibiting lower affinities (*h*DMX_335-349_^pSer342^, *h*DM2_160-171_^pSer166^ and *h*DM2_180-192_^pSer186^) it was only possible to determine steady-state affinities, but not their kinetic profile (see supporting information). The doubly phosphorylated peptides showed the expected increase in affinity, which could be attributed to a much slower k_off_ rate in comparison to singly phosphorylated *h*DMX_361-374_^pSer367^ and *h*DMX_335-349_^pSer342^ peptides.

The 1:1 stoichiometry between the doubly phosphorylated peptides and 14-3-3 contrasts with previously reported work,^58-59^ although it may be that the length and/or accessible conformation of the *h*DMX and *h*DM2 peptides is incompatible with simultaneous engagement of both 14-3-3 protomers in the dimeric structure. To further explore this behaviour and understand the basis of cooperativity and affinity enhancement we performed additional experiments. To exclude possible formation of higher order oligomers (e.g. 14-3-3 tetramers held together by two peptides bridging two dimers), an analytical ultracentrifugation experiment was performed. 14-3-3η (at the concentration of 14 μM, approximately corresponding to the second binding events observed in FA, Fig 2c) and *h*DMX_335-373_^pSer342/pSer367^ peptide in two different concentration ratios (1:1 and 1:0.5, protein versus peptide) were analysed. The sedimentation coefficient value remained unchanged for 14-3-3 protein alone and when varying peptide/protein concentrations, indicating that only one dimer of 14-3-3η is involved in binding (see supporting information Fig. S10). Secondly, peptide length may also have a significant effect on the affinity and binding mode. This has been demonstrated in a recent study with p53 peptides with 14-3-3σ, where variation in the peptide length (from 9 to 32 residues) correlated with increased binding affinities.^61^ Two long variants of *h*DMX_335-373_^pSer342/pSer367^ peptide were synthesized, with only one phosphorylation site: pSer367 or pSer342, extending the peptide length by 26 amino acids. The two peptides were tested in FA assays to decouple the influence of peptide length from the co-operative influence of having two phosphosites. The *h*DMX_335-373_^pSer342^ peptide showed an increase in binding affinity with 14-3-3η (*K*_d_= 5.4 ± 1.1 μM) as expected, in comparison to the short version of the *h*DMX_335-349_^pSer342^ peptide (*K*_d_ = 20.0 ± 1.2 μM). On the other hand, increasing the peptide length for *h*DMX_335-373_^pSer367^ decreased the binding affinity (*K*_d_ = 0.8 ± 0.1 μM) and significantly lowered the maximum anisotropy signal. The lower max. signal observed might be due to the fluorophore being moved approximately 20 amino acids further from the binding pSer-367 (see earlier discussion on fluorophore mobility). Whilst there is a moderate increase in binding affinity of *h*DMX_335-373_^pSer342^ on increasing the peptide length this is insufficient for high affinity interaction. Similarly, the slight loss in potency for the longer *h*DMX_335-373_^pSer367^ peptide is noteworthy in considering the interaction with full length protein as these data hint at interference from the additional residues. Taken together, the data indicate that proximal phosphorylation of both sites in *h*DMX_335-373_^pSer342/pSer367^ is required for high affinity 1:1 stoichiometric binding between *h*DMX and 14-3-3 protomers. Given the absence of 1:2 binding, the presence of multiple sites may increase affinity through statistical rebinding or interaction with a second non-canonical phosphate binding site on the surface of 14-3-3. The second binding event observed only in fluorescence anisotropy experiments, may represent a further non-specific binding event or an interaction linked to 14-3-3 dimerization^57^ and will be investigated in future work.

### Structural characterization of the interaction between 14-3-3 and *h*DMX or *h*DM2 peptides

The W shaped structure of a 14-3-3 dimer provides a groove for phosphorylated client proteins to bind. One dimer usually accommodates two copies of phosphorylated sites from one target, but there are examples of two different targets simultaneously binding to one dimer of 14-3-3 proteins.^58-59^ Phosphorylated Ser/Thr residues from partner proteins generally bind directly between Lys49 and Arg56 in helix α3, and Arg127 and Tyr128 in helix α5 as they form a basic binding pocket in otherwise acidic surroundings. Structural data were obtained by solving three crystal structures of *h*DMX_361-374_^pSer367^ (PDB: 6YR5), *h*DM2_180-192_^pSer186^ (PDB: 6YR6) and *h*DMX_335-373_^pSer342/pSer367^ (PDB: 6YR7) in complex with 14-3-3σΔC (the C-terminus was truncated for the crystallography purposes). Refinement statistics for all three structures are given in the ESI as are additional images of each structure with electron density for peptides in each monomer of 14-3-3σ.

The *h*DMX_361-374_^pSer367^/14-3-3σ structure was solved to a resolution of 2.25 Å with a dimer and two additional monomers of 14-3-3σ (in green and blue) in the asymmetric unit, each binding a *h*DMX_361-374_^pSer367^ peptide (cyan) in a conserved amphipathic groove (Fig. 4a and supporting information Fig S11 for additional images). Ten amino acids from *h*DMX_361-374_^pSer367^ (RTI(pS) APVVRP) could be built into the electron density map. The phosphoserine of *h*DMX_361-374_^pSer367^ peptide is positioned in a basic pocket of 14-3-3σ making polar contacts with Lys49, Arg56, Tyr119 and Arg129 (presented as dashed lines), whilst Asn226 makes polar contacts with the backbone of the peptide (Ile-1 with reference to pSer-367), in agreement with previously published 14-3-3 structures.^37^ Additional polar contacts were found between Arg-3 of the *h*DMX_361-374_^pSer367^ and Glu182 of 14-3-3σ, alongside Ala+1 and Thr+2 on the peptide backbone with Asn175 and Trp230 on the protein respectively. Hydrophobic contacts were found between Ile-1, Ala+1 and Pro+2 of the peptide with multiple residues on the protein (presented as spheres).

**Figure 4.**
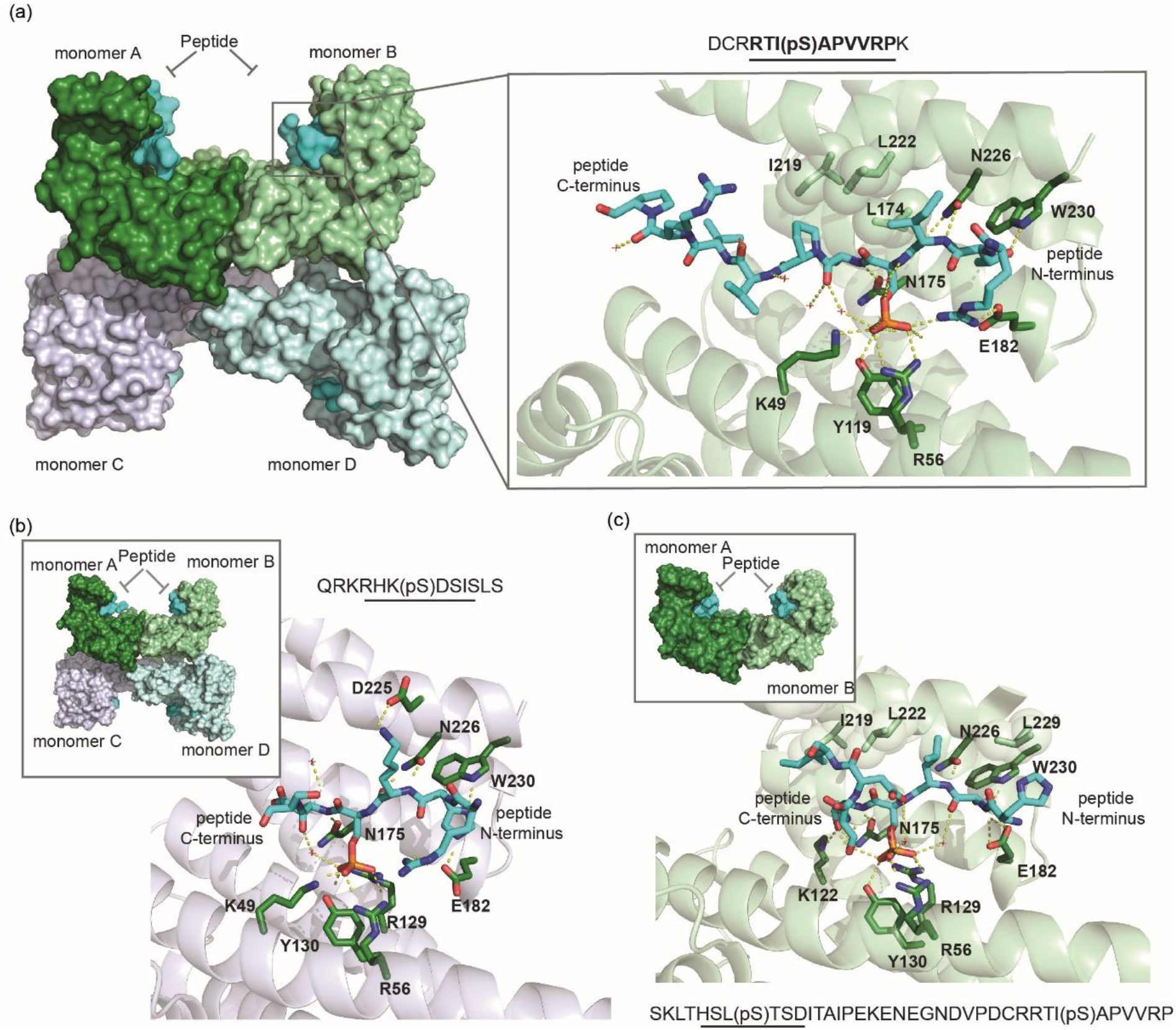
Interactions between 14-3-3 and hDM peptides (a) *h*DMX_361-374_^pSer367^/14-3-3σ structure (PDB: 6YR5); left hand side shows assembly of the *h*DMX_361-374_^pSer367^/14-3-3σ complexes in the asymmetric unit with each singly phosphorylated peptide (cyan surface) bound to a 14-3-3σ monomer (dark green, light green, light blue or light lilac surface), in its conserved amphipathic groove, right hand side shows monomer B (light green fold, 14-3-3σ side chains shown as sticks with carbon dark green, nitrogen dark blue and oxygen red, *h*DMX_361-374_^pSer367^ shown as sticks, carbon in cyan, phosphorous orange, nitrogen and oxygen as above, non-covalent contacts shown as dashed lines) (b) *h*DM2_180-192_^pSer186^/14-3-3σ (PDB: 6YR6) monomer C (light lilac fold, 14-3-3σ and *h*DM2_180-192_^pSer186^ side chains shown as sticks with colour coding as above), inset shows complex in asymmetric unit (colour coding as above) (c) *h*DMX_335-373_^pSer342/pSer367^/14-3-3σ structure (PDB: 6YR7) monomer B (light green fold, 14-3-3σ and *h*DMX_335-373_^pSer342/pSer367^ side chains shown as sticks with colour coding as above), inset shows complex in asymmetric unit (colour coding as above)

Structural evidence for interaction of *h*DM2 with 14-3-3 proteins was obtained by solving the novel crystal structure of the *h*DM2_180-192_^pSer186^ peptide in complex with 14-3-3σ, to a resolution of 1.75 Å. The complex crystallized in space group *P*1 with the asymmetric unit containing one dimer along with an additional two monomers of 14-3-3σ (monomers A-D) with pSer186 peptide in each groove (in cyan). Six amino acids of the peptide were resolved in electron density: RHK(pS)DS (Fig. 4b see supporting information Fig S12 for additional images). pSer of the peptide is always located in the same position, between Lys49, Arg56, Arg129 and Tyr130 and the peptide is stabilized by polar contacts between Lys-1 and Asp225, His-2 and Glu182, Trp230, and the peptide backbone with Asn175 and Asn226. Arg-3 and Ser+2 peptide residues are also stabilized through hydrogen bonds with water molecules.

The structure of a doubly phosphorylated peptide *h*DMX_335-373_^pSer342/pSer367^ in complex with 14-3-3σ was solved to 2.1 Å resolution (Fig. 4c). Here 14-3-3σ crystallized as a dimer in space group C121 with the asymmetric unit containing one dimer (green) with additional electron density for the peptide (cyan) observed in both monomeric units of 14-3-3σ (Fig. 4c and supporting information Fig. S13 for additional images). Based on previously published structures of doubly phosphorylated peptides, it was expected to observe pSer342 binding to monomer B and the pSer367 binding to monomer A.^31, 58, 62^ The electron density within the binding groove of monomer B could unambiguously be assigned to the pSer342, whereby eight amino acids of the peptide could be resolved (HSL(pS)TSDI). The peptide was observed to adopt an unusual conformation by taking a sharp turn. A closer examination of the pSer342 binding site in monomer B of 14-3-3σ shows pSer binding to a basic pocket as expected, making polar contacts with Arg56, Arg129 and Tyr130. In this orientation, the Ile+4 from the peptide makes hydrophobic contacts with Leu218 and Ile219 on the protein (represented as spheres). Peptide residues Thr+1 and Ser+2 make polar contacts with Lys122, Asn175 and Glu182, Trp230, while Asn226 makes polar contacts with the backbone of the peptide. Overall, the C-terminus of the peptide takes a sharp turn after the pSer-Thr residues and this peptide conformation is stabilized by hydrogen bonds between water molecules and Thr+1, Ser+2 and Asp+3 peptide residues. In contrast the electron density map within the binding groove of monomer A indicated partial ligand occupancy and could be interpreted in different ways, indicating an overlay of multiple conformations (both pSer342 and pSer367 sites, with the dominant electron density map ascribable to pSer342 binding see supporting information Fig S13 for additional images and discussion).

## Discussion

We used FA, ITC and SPR to determine the binding affinity of *h*DMX and *h*DM2 peptides to 14-3-3 proteins. *h*DMX_335-349_^pSer342^, *h*DMX_361-374_^pSer367^, *h*DM2_160-171_^pSer166^ and *h*DM2_180-192_^pSer186^ mimicking one phosphorylation site fit well to a 1:1 binding isotherm and exhibited greatest affinity towards the 14-3-3η isoform, followed by γ, β, τ, ζ, σ and ε, respectively. The highest affinity peptide (*h*DMX_361-374_^pSer367^) bound strongly (*K*_d_ = 98.8 ± 4.6 nM), whereas the remaining sequences bound 14-3-3 less tightly (*h*DMX_335-349_^pSer342^ *K*_d_ = 20.0 ± 1.2 μM {14-3-3η isoform}, *h*DM2_160-171_^pSer166^ *K*_d_ = 10.2 ± 0.3 μM and *h*DM2_180-192_^pSer186^ *K*_d_ = 31.6 ± 1.9 μM). Interestingly, different maximum anisotropies were observed for peptide binding across the 14-3-3 isoforms, which may reflect a difference in the 14-3-3 dimerization affinity^57^ or mobility of the fluorophore. Binding experiments with doubly phosphorylated peptides were observed to result in two binding events. As expected for a multivalent interaction with 14-3-3 proteins,^60^ the first binding event in both cases was higher affinity than either monovalent interaction, whereas the second binding event was in the micromolar regime (for 14-3-3η, *h*DMX_335-373_^pSer342/pSer367^ *K*_d1_ = 30.3 ± 6.4 nM and *K*_d2_ = 10.67 ± 1.2 μM, *h*DM2_160-192_^pSer166/pSer186^; *K*_d1_ = 49.6 ± 12.1 nM and *K*_d2_ = 17.5 ± 2.5 μM). This enhancement in affinity for the first binding event was also observed in ITC experiments which lacked the second binding event (but were performed by titration of peptide into protein in contrast to the FA experiments) and SPR experiments which indicated that a slower *k*_off_ was associated with co-operative binding of doubly phosphorylated peptides. Importantly, for the doubly phosphorylated peptides, binding was negatively co-operative (i.e. less than additive (Table 4).

**Table 4.** Thermodynamic values for binding of *h*DM2 and *h*DMX peptides to 14-3-3η

| Peptide | $\Delta G_{14-3-3\eta}$ (kJ/mol) |
| --- | --- |
| *hDMX*<sub>335-349</sub><sup>pSer342</sup> | 21.1 |
| *hDMX*<sub>361-374</sub><sup>pSer367</sup> | 34.3 |
| *hDMX*<sub>335-373</sub><sup>pSer342/pSer367</sup> | 37.2 |
| *hDM2*<sub>160-171</sub><sup>pSer166</sup> | 22.8 |
| *hDM2*<sub>180-192</sub><sup>pSer186</sup> | 20.0 |
| *hDM2*<sub>160-192</sub><sup>pSer166/pSer186</sup> | 36.0 |

Taken together the data can be rationalized by assuming that the dimerization affinity of 14-3-3 differs across isoforms and occurs within the concentration regime of the FA experiments (Fig. 5). As the titration progresses, once 1:1 stoichiometry is attained, further addition of 14-3-3 results in unbound 14-3-3 monomer and eventually all peptide will be bound in complexes containing one peptide to two 14-3-3 monomers. For the doubly phosphorylated peptides early in the titration peptides bind with high affinity in 1:1 stoichiometry – higher affinity likely results from statistical rebinding or a second phosphosite on the surface of the 14-3-3 protein. The second binding event either derives from a non-specific interaction, or arises from 14-3-3 dimerization^57^ but which is visible in the titration as a distinct binding event due to the strong affinity of the first event. Such behaviour is not observed in ITC experiments because peptide is titrated into 14-3-3 protein so its concentration does not vary, and, peptide becomes the saturating component. Considering the structural data for *h*DMX_361-374_^pSer367^, *h*DM2_180-192_^pSer186^ and *h*DMX_335-373_^pSer342/pSer367^ bind to 14-3-3 directly between K49 and R56 in helix α3, and R127 and Y128 in helix α5 as is typical for 14-3-3 client proteins. The *h*DMX_361-374_^pSer367^peptide is a Mode I sequence (RXXpS/pTXP) which can account for its enhanced affinity in comparison to the other sequences.^37, 54^ Curiously, for the structure containing the longer *h*DMX_335-373_^pSer342/pSer367^ peptide, the sequence surrounding pSer342 dominates in the electron density for bound ligand despite being the weaker 14-3-3 binding site of the two individual pSer sequences, this may favour the hypothesis of a second phosphorecognition site over the statistical rebinding model. Moreover, in the context of earlier cellular studies^26, 39-43, 46-50 51^ the data highlight the complex nature by which phosphorylation regulates *h*DMX in particular.^26, 39-43^ pSer367 may be sufficient for 14-3-3 binding and the consequent effects on localization, stability and ultimately p53 function; the presence of both pSer342 and pSer367 confers some increase in 14-3-3 binding affinity (particularly for other isoforms). In contrast, a much clearer difference in binding affinity for *h*DM2_160-192_^pSer166/pSer186^ is observed in contrast to both monophosphorylated peptides and points to a functional requirement for double phosphorylation.

**Figure 5.**
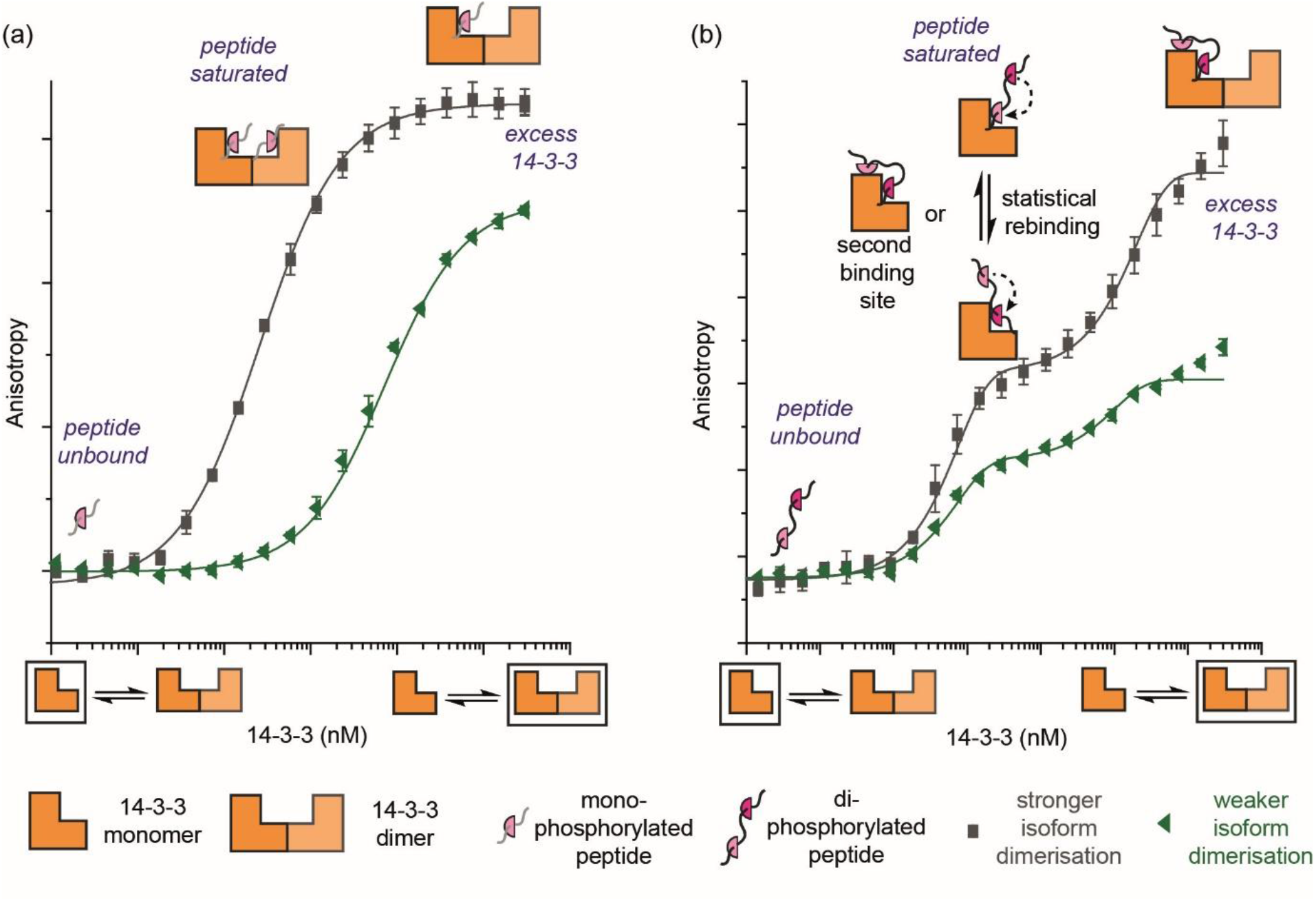
Schematic overview depicting equilibrium behaviour and potential hypotheses for isoform dependent differences in anisotropies linked to 14-3-3 dimerization alongside reasons for enhanced affinity of doubly phosphorylated peptides.

## Conclusions

We have performed a biophysical and structural study on the interaction of 14-3-3 with peptides derived from *h*DMX and *h*DM2. We have shown that peptides derived from *h*DMX containing pSer342 or pSer367 and from *h*DM2 containing pSer166 or pSer186 exhibit affinity for 14-3-3 as demonstrated by FA, ITC and SPR. Structural analyses reveal that the peptides bind to 14-3-3 directly between K49 and R56 in helix α3, and R127 and Y128 in helix α5 as is typical for 14-3-3 client proteins. For high affinity binding, proximal phosphosites are beneficial and in the case of *h*DM2 essential. In contrast to other doubly phosphorylated peptides of this length the interaction involves one peptide for each 14-3-3 protomer.^58-60^ The additional affinity likely derives from a reduced *k*_off_ rate for doubly phosphorylated peptides. Importantly, the results suggest that sequential phosphorylation of proximal sites in both *h*DMX and *h*DM2 plays a role in promoting changes in their localization and degradation in a cellular context. Thus, these data provide structural insight on the indirect affects of 14-3-3 on the p53 pathway, which may inform the development of chemical intervention strategies (e.g. use of kinase inhibitors) and provide starting points for structure based ligand design of 14-3-3/*h*DMX and 14-3-3/*h*DM2 modulators, both stabilizers and inhibitors.

## Acknowledgements

We would like to thank Dr Iain Manfield for his support with ITC and SPR measurements and ongoing collaboration with AstraZeneca.

## Author contribution statement

SS, CO, SLW and AJW conceived and designed the research program, SS designed studies and performed the research with support from MW, CT and CO on crystallographic analyses. The manuscript was written by SS and AJW and edited into its final form by SLW and AJW with contributions from all authors.

