## Supporting Information for "Understanding the interaction of 14-3-3 proteins with *h*DMX and *h*DM2: a structural and biophysical study"

### Additional Figures

#### Protein Sequences and Alignment

```
P27348 1433T_HUMAN      1   --MEKTELIQKAKLAEQAERYDDMATCMKAVTEQGAELSNEERNLLSVAYKNVVGGRRSA    58
P31946 1433B_HUMAN      1   MTMDKSELVQKAKLAEQAERYDDMAAAMKAVTEQGHELSNEERNLLSVAYKNVVGARRSS    60
P61981 1433G_HUMAN      1   -MVDREQLVQKARLAEQAERYDDMAAAMKNVTELNEPLSNEERNLLSVAYKNVVGARRSS    59
P62258 1433E_HUMAN      1   -MDDREDLVYQAKLAEQAERYDEMVESMKKVAGMDVELTVEERNLLSVAYKNVIGARRAS    59
P63104 1433Z_HUMAN      1   --MDKNELVQKAKLAEQAERYDDMAACMKSVTEQGAELSNEERNLLSVAYKNVVGARRSS    58
Q04917 1433F_HUMAN      1   -MGDREQLLQRARLAEQAERYDDMASAMKAVTELNEPLSNEDRNLLSVAYKNVVGARRSS    59
P31947 1433S_HUMAN      1   --MERASLIQKAKLAEQAERYEDMAAFMKGAVEKGEELSCEERNLLSVAYKNVVGGQRAA    58
                               :: .*: :*:********::*.  **  ..   .   *: *:***********:*.:*::

P27348 1433T_HUMAN     59   WRVISSIEQKT--DTSDKKLQLIKDYREKVESELRSICTTVLELLDKYLIANAT--NPES   114
P31946 1433B_HUMAN     61   WRVISSIEQKT--ERNEKKQQMGKEYREKIEAELQDICNDVLELLDKYLIPNAT--QPES   116
P61981 1433G_HUMAN     60   WRVISSIEQKTSADGNEKKIEMVRAYREKIEKELEAVCQDVLSLLDNYLIKNCSETQYES   119
P62258 1433E_HUMAN     60   WRIISSIEQKEENKGGEDKLKMIREYRQMVETELKLICCDILDVLDKHLIPAAN--TGES   117
P63104 1433Z_HUMAN     59   WRVVSSIEQKT--EGAEKKQQMAREYREKIETELRDICNDVLSLLEKFLIPNAS--QAES   114
Q04917 1433F_HUMAN     60   WRVISSIEQKTMADGNEKKLEKVKAYREKIEKELETVCNDVLSLLDKFLIKNCNDFQYES   119
P31947 1433S_HUMAN     59   WRVLSSIEQKSNEEGSEEKGPEVREYREKVETELQGVCDTVLGLLDSHLIKEAG--DAES   116
                            **::******    .   :.*    : **: :* **. :*   :* :*:..**  .     **

P27348 1433T_HUMAN    115   KVFYLKMKGDYFRYLAEVACGDDRKQTIDNSQGAYQEAFDISKKEMQPTHPIRLGLALNF   174
P31946 1433B_HUMAN    117   KVFYLKMKGDYFRYLSEVASGDNKQTTVSNSQQAYQEAFEISKKEMQPTHPIRLGLALNF   176
P61981 1433G_HUMAN    120   KVFYLKMKGDYYRYLAEVATGEKRATVVESSEKAYSEAHEISKEHMQPTHPIRLGLALNY   179
P62258 1433E_HUMAN    118   KVFYYKMKGDYHRYLAEFATGNDRKEAAENSLVAYKAASDIAMTELPPTHPIRLGLALNF   177
P63104 1433Z_HUMAN    115   KVFYLKMKGDYYRYLAEVAAGDDKKGIVDQSQQAYQEAFEISKKEMQPTHPIRLGLALNF   174
Q04917 1433F_HUMAN    120   KVFYLKMKGDYYRYLAEVASGEKKNSVVEASEAAYKEAFEISKEQMQPTHPIRLGLALNF   179
P31947 1433S_HUMAN    117   RVFYLKMKGDYYRYLAEVATGDDKKRIIDSARSAYQEAMDISKKEMPPTNPIRLGLALNF   176
                            :*** ******.***:*.* *:.:    . :  **. * :*:  .: **:*********:

P27348 1433T_HUMAN    175   SVFYYEILNNPELACTLAKTAFDEAIAELDTLNEDSYKDSTLIMQLLRDNLTLWTSDSAG   234
P31946 1433B_HUMAN    177   SVFYYEILNSPEKACSLAKTAFDEAIAELDTLNEESYKDSTLIMQLLRDNLTLWTSENQG   236
P61981 1433G_HUMAN    180   SVFYYEIQNAPEQACHLAKTAFDDAIAELDTLNEDSYKDSTLIMQLLRDNLTLWTSDQQD   239
P62258 1433E_HUMAN    178   SVFYYEILNSPDRACRLAKAAFDDAIAELDTLSEESYKDSTLIMQLLRDNLTLWTSDMQG   237
P63104 1433Z_HUMAN    175   SVFYYEILNSPEKACSLAKTAFDEAIAELDTLSEESYKDSTLIMQLLRDNLTLWTSDTQG   234
Q04917 1433F_HUMAN    180   SVFYYEIQNAPEQACLLAKQAFDDAIAELDTLNEDSYKDSTLIMQLLRDNLTLWTSDQQD   239
P31947 1433S_HUMAN    177   SVFHYEIANSPEEAISLAKTTFDEAMADLHTLSEDSYKDSTLIMQLLRDNLTLWTADNAG   236
                            ***:*** * *: *  *** :**:*:*:*.**.*:*******************::  .

P27348 1433T_HUMAN    235   EECDAAEGAEN-------                                           245
P31946 1433B_HUMAN    237   DEGDAGEGEN--------                                           246
P61981 1433G_HUMAN    240   DDGGEGNN----------                                           247
P62258 1433E_HUMAN    238   DGEEQNKEALQDVEDENQ                                           255
P63104 1433Z_HUMAN    235   DEAEAGEGGEN-------                                           245
Q04917 1433F_HUMAN    240   EEAGEGN-----------                                           246
P31947 1433S_HUMAN    237   EEGGEAPQEPQS------                                           248
                            :
```

**Figure S1. Sequence alignment of** 14-3-3 Proteins. UniProt codes for seven human 14-3-3 isoforms: τ- P27348, η- Q04917, σ- P31947, ε-P62258, β- P31946, γ- P61981, ζ- P63104 (conserved amino acids are in dark grey whilst similar amino acids are in light grey).

UniProt code: O15151,*h*DMX_HUMAN Protein

```
MTSFSTSAQCSTSDSACRISPGQINQVRPKLPLLKILHAAGAQGEMFTVKEVMHYLGQYI
MVKQLYDQQEQHMVYCGGDLLGELLGRQSFSVKDPSPLYDMLRKNLVTLATATTDAAQTL
ALAQDHSMDIPSQDQLKQSAEESSTSRKRTTEDDIPTLPTSEHKCIHSREDEDLIENLAQ
DETSRLDLGFEEWDVAGLPWWFLGNLRSNYTPRSNGSTDLQTNQDVGTAIVSDTTDDLWF
LNESVSEQLGVGIKVEAADTEQTSEEVGKVSDKKVIEVGKNDDLEDSKSLSDDTDVEVTS
EDEWQCTECKKFNSPSKRYCFRCWALRKDWYSDCSKLTHSLSTSDITAIPEKENEGNDVP
DCRRTISAPVVRPKDAYIKKENSKLFDPCNSVEFLDLAHSSESQETISSMGEQLDNLSEQ
RTDTENMEDCQNLLKPCSLCEKRPRDGNIIHGRTGHLVTCFHCARRLKKAGASCPICKKE
IQLVIKVFIA
```

UniProt code: Q00987, *h*DM2_HUMAN E3 ubiquitin-protein ligase

```
MCNTNMSVPTDGAVTTSQIPASEQETLVRPKPLLLKLLKSVGAQKDTYTMKEVLFYLGQY
IMTKRLYDEKQQHIVYCSNDLLGDLFGVPSFSVKEHRKIYTMIYRNLVVVNQQESSDSGT
SVSENRCHLEGGSDQKDLVQELQEEKPSSSHLVSRPSTSSRRRAISETEENSDELSGERQ
RKRHKSDSISLSFDESLALCVIREICCERSSSSESTGTPSNPDLDAGVSEHSGDWLDQDS
VSDQFSVEFEVESLDSEDYSLSEEGQELSDEDDEVYQVTVYQAGESDTDSFEEDPEISLA
DYWKCTSCNEMNPPLPSHCNRCWALRENWLPEDKGKDKGEISEKAKLENSTQAEEGFDVP
DCKKTIVNDSRESCVEENDDKITQASQSQESEDYSQPSTSSSIIYSSQEDVKEFEREETQ
DKEESVESSLPLNAIEPCVICQGRPKNGCIVHGKTGHLMACFTCAKKLKKRNKPCPVCRQ
PIQMIVLTYFP
```

**Figure S2** *h*DMX and *h*DM2 sequences highlighting serine and threonine residues bold and blue).

```
Q00987 MDM2_HUMAN          1   MCNTNMSVPTDGAVTTSQIPASEQETLVRPKPLLLKLLKSVGAQKDTYTMKEVLFYLGQY
O15151 MDM4_HUMAN          1   MTSFSTSAQCSTSDS-ACRISPGQINQVRPKLPLLKILHAAGAQGEMFTVKEVMHYLGQY
                               *  . .  *.   . : : :    :    *  . ****   ***:*::.***  :  :*:***:.*****

Q00987 MDM2_HUMAN         61   IMTKRLYDEKQQHIVYCSNDLLGDLFGVPSFSVKEHRKIYTMIYRNLVVVNQQESSDSGT
O15151 MDM4_HUMAN         60   IMVKQLYDQQEQHMVYCGGDLLGELLGRQSFSVKDPSPLYDMLRKNLVTLATATTDAAQT
                               **.*:***:::**:***..****:*:*   *****:    :* *: :***.:      :.  : *

Q00987 MDM2_HUMAN        121   SVSENRCHLEGGSDQKDLVQELQEEKPS-----SSHLVSRPSTSSRRRAISETEENSDEL
O15151 MDM4_HUMAN        120   LALA-QDHSMDIPSQDQLKQSAEESSTSRKRTTEDDIPTLPTS--EHKCIHS-REDEDLI
                                .    : *   .   .*.:* *.  :*..  *        ...: : *::   .::.*  .  .*:.* :

Q00987 MDM2_HUMAN        176   SGERQRKRHKSDSISLSFDES----LALCVI---REICCERSSSSESTGTPSNPDLDAG-
O15151 MDM4_HUMAN        176   ENLA---QDETSRLDLGFEEWDVAGLPWWFLGNLRSNYTPRSNG--STDLQTNQDVGTAI
                               ..       :.::. :.*.*:*       *    .:     *.      **..   **.   :* *:.:.

Q00987 MDM2_HUMAN        228   -VSEHSGDWLDQDSVSDQFSVEFEVESLDSEDYSLSEEGQELSDEDDEVYQVTVYQAGE-
O15151 MDM4_HUMAN        231   VSDTTDDLWFLNESVSEQLGVGIKVEAADTEQTSEE--V-------GKVSDKKVIEVGKN
                                 .   .. *:  ::***:*:.*  ::**:  *:*:  *  .             .:*  :  .*  :.*:

Q00987 MDM2_HUMAN        286   -----SDTDSFEEDPEISLADYWKCTSCNEMNPPLPSHCNRCWALRENWLPEDKGKDKGE
O15151 MDM4_HUMAN        282   DDLEDSKSLSDDTDVEVTSEDEWQCTECKKFNSPSKRYCFRCWALRKDWYSDCSKLTHS-
                                    *.:  *  :  *  *::   *  *:**.*:::* *    :*  ******::*   :  .    :.

Q00987 MDM2_HUMAN        341   ISEKAKLENSTQAEEGFDVPDCKKTIVNDSR---ESCVEENDDKITQASQSQESEDYSQP
O15151 MDM4_HUMAN        341   LSTSDITAIPEKENEGNDVPDCRRTISAPVVRPKDAYIKKENSKLFDPCNSVEFLDLAHS
                               :*  .          :  :**  *****::**           ::  :::::.*:  :  .:*  *   *  ::

Q00987 MDM2_HUMAN        398   STSSSIIYSSQEDVKEFEREETQDKEESVESSLPLNAIEPCVICQGRPKNGCIVHGKTGH
O15151 MDM4_HUMAN        401   SESQETISSMGEQLDNLSEQR----TDTENMEDCQNLLKPCSLCEKRPRDGNIIHGRTGH
                               *  *..  *  *   *::.::..:.         ::  :  .     *  ::**  :*:  **::*  *:**:***

Q00987 MDM2_HUMAN        458   LMACFTCAKKLKKRNKPCPVCRQPIQMIVLTYFP
O15151 MDM4_HUMAN        457   LVTCFHCARRLKKAGASCPICKKEIQLVIKVFIA
                               *::**  **::***  .   **:*::  **:::  .::
```

**Figure S3**. Sequence alignment for *h*DM2 and *h*DMX proteins (conserved amino acids are in dark grey whilst similar amino acids are in light grey).

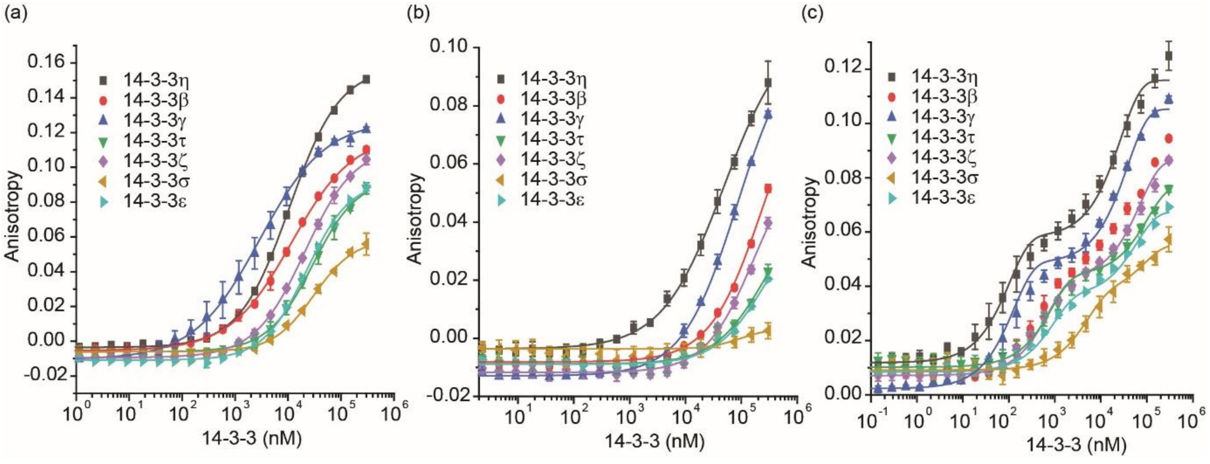

**Figure S4**. Fluorescence anisotropy assays for the *h*DM2 peptides. (a) *h*DM2$_{160\text{-}171}^{pSer166}$ and *h*DM2$_{180\text{-}192}^{pSer186}$ (c) *h*DM2$_{160\text{-}192}^{pSer166/pSer186}$ with all isoforms of 14 3 3 (FAM tracer peptide 50 nM, 0.1 nM - 300 µM protein, in 10 mM HEPES, 150 mM NaCl, 0.1% Tween 20, 0.1% BSA, pH 7.4, concentration of proteins given as 14-3-3 monomer concentration)

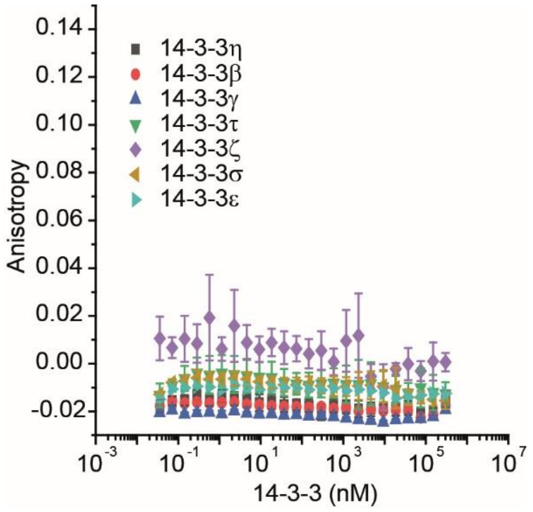

**Figure S5**. Fluorescence anisotropy assays for the *h*DMX$_{144\text{-}158}^{pThr151}$ peptide with all isoforms of 14-3-3 (FAM tracer peptide 50 nM, 0.1 nM - 300 µM protein, in 10 mM HEPES, 150 mM NaCl, 0.1% Tween 20, 0.1% BSA, pH 7.4, concentration of proteins given as 14-3-3 monomer concentration)

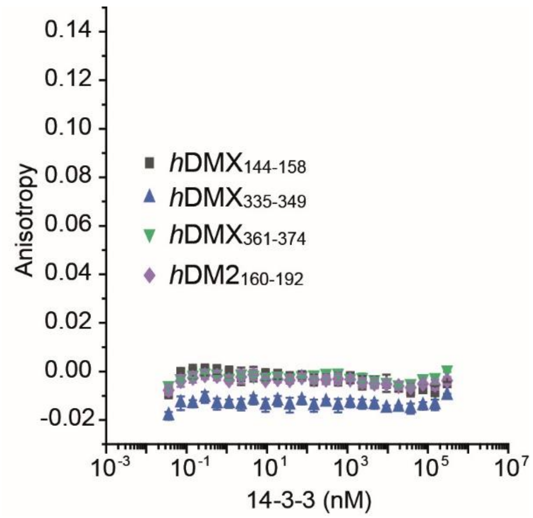

**Figure S6**. Fluorescence anisotropy assays for unphosphorylated *h*DMX and *h*DM2 peptides with 14-3-3η (FAM tracer peptide 50 nM, 0.1 nM - 300 µM protein, in 10 mM HEPES, 150 mM NaCl, 0.1% Tween 20, 0.1% BSA, pH 7.4, concentration of proteins given as 14-3-3 monomer concentration)

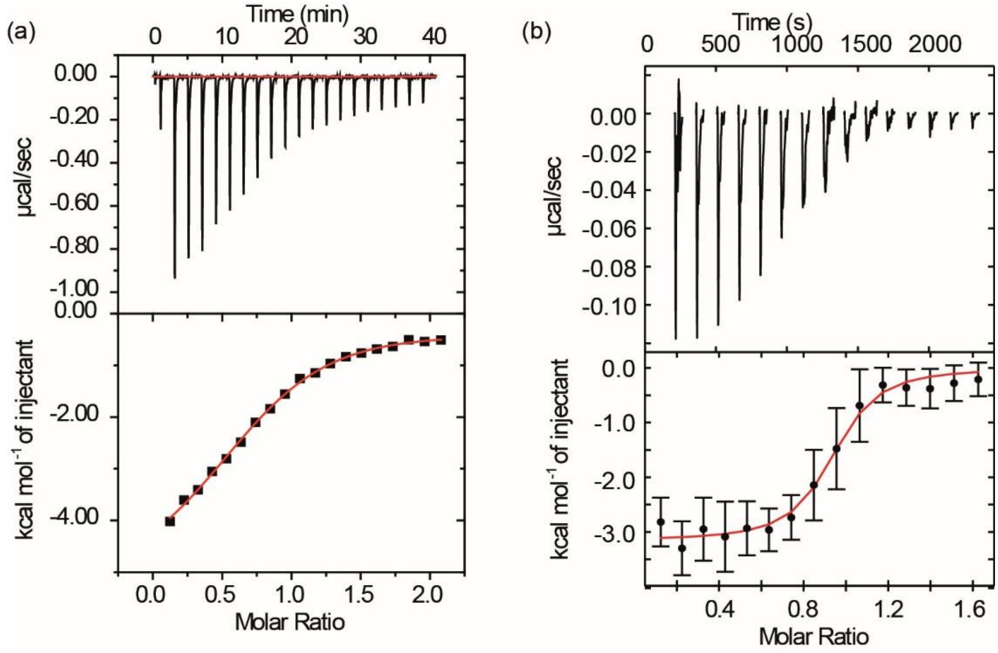

**Figure S7**. ITC data for the interaction of *h*DMX and *h*DM2 peptides with 14-3-3η; (a) *h*DMX$_{335\text{-}349}^{pSer342}$; (b) *h*DM2$_{160\text{-}192}^{pSer166/pSer186}$ fitted with SEDPHAT (peptides were titrated into 14-3-3 η {0.1M for mono phosphorylated peptide and 0.02M for doubly phosphorylated peptide}, 25°C, 25 mM HEPES pH 7.5, 100 mM NaCl, 10 mM $MgCl_2$, 0.5 mM TCEP).

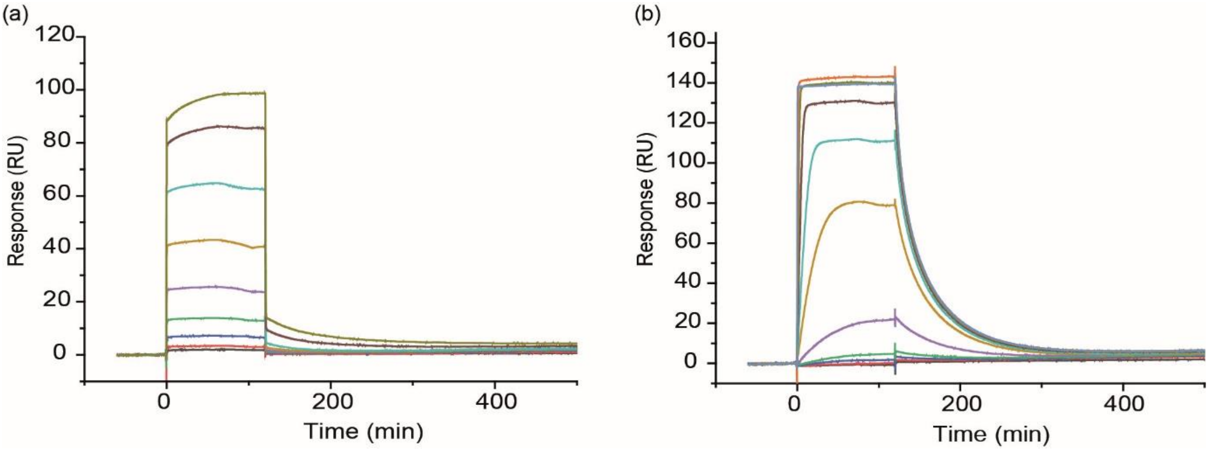

**Figure S8**. Dose response SPR experiments for the binding of *h*DMX and *h*DM2 peptides to immobilized 14-3-3η; (a) *h*DMX$_{335\text{-}349}^{\text{pSer342}}$; (b) *h*DM2$_{160\text{-}192}^{\text{pSer166/pSer186}}$ (peptides – concentration at 10x the $K_d$ for each peptide – were passed over immobilized 14-3-3η, 25°C, 25 mM HEPES pH 7.5, 100 mM NaCl, 10 mM $MgCl_2$; experiments were performed in a multicycle kinetic format and data was fitted to a Langmuir model. $K_d$ values were determined by fitting maximal response level at the end of injection against protein concentration using a steady state affinity model in the Biocore evaluation software.

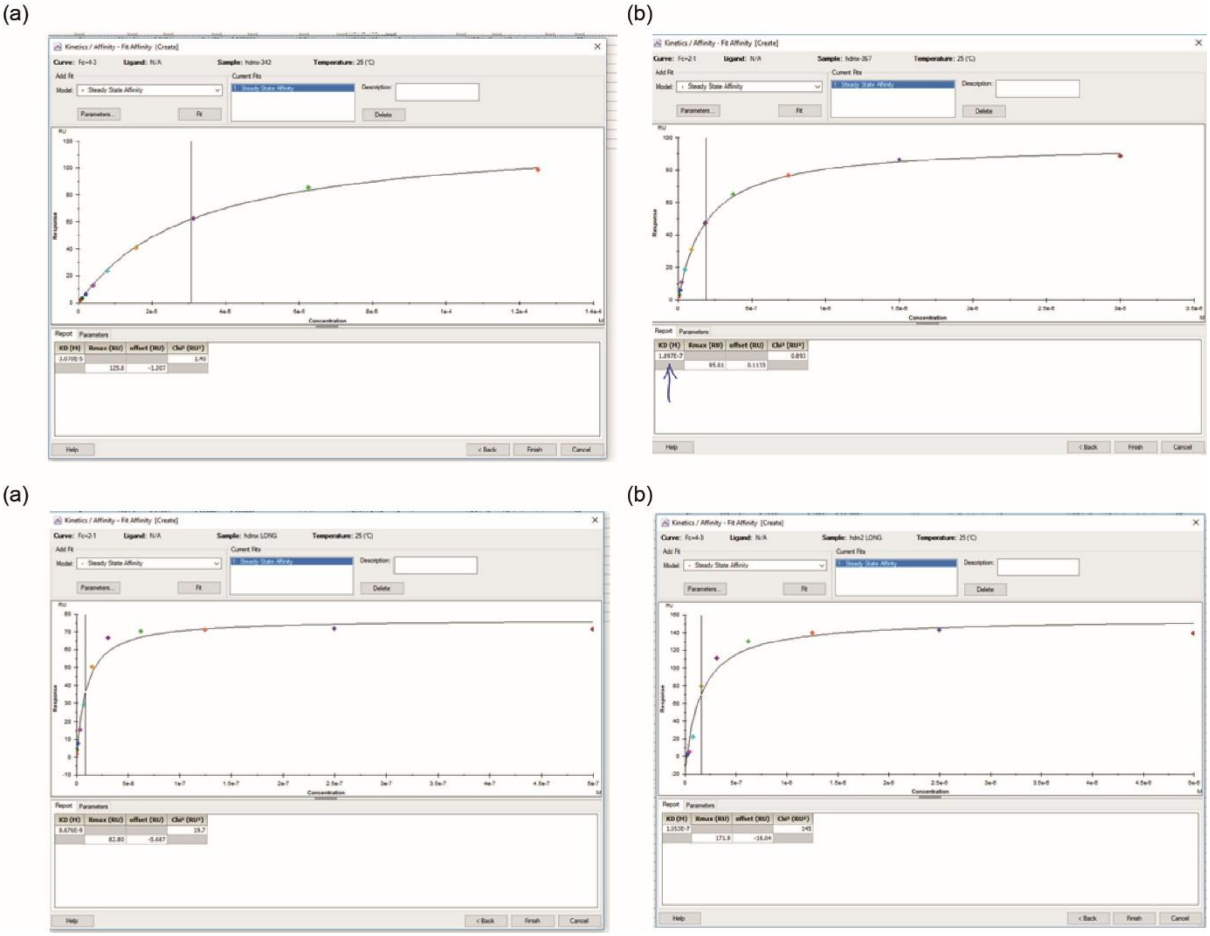

**Figure S9**. Data Fitting for Dose response SPR experiments for the binding of *h*DMX and *h*DM2 peptides to immobilized 14-3-3η; (a) *h*DMX$_{335\text{-}349}^{\text{pSer342}}$; (b) *h*DMX$_{361\text{-}374}^{\text{pSer367}}$, (c) *h*DMX$_{335\text{-}373}^{\text{pSer342/pSer367}}$ and (d) *h*DM2$_{160\text{-}192}^{\text{pSer166/pSer186}}$.

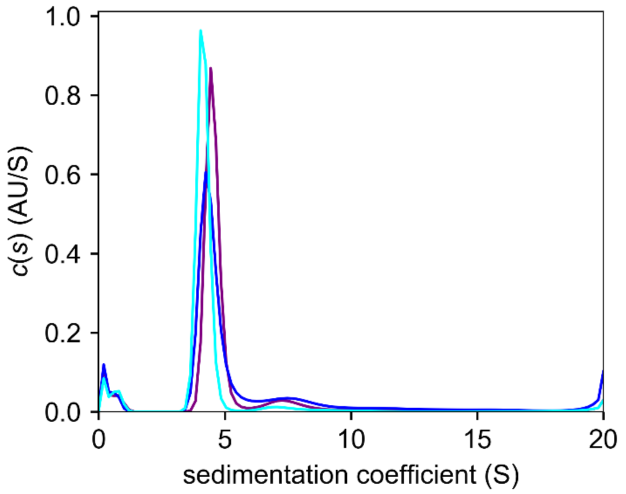

**Figure S10**. AUC data for 14-3-3η alone (purple) and in a mixture with *h*DMX$_{335\text{-}374}^{pSer342/pSer367}$ (0.5 eq dark blue, 1.0 eq. light blue), indicating only one physiological dimer is involved in the binding (14 μM of protein used with two different rations of *h*DMX$_{335\text{-}374}^{pSer342/pSer367}$, in 10 mM HEPES, 150 mM NaCl, 0.1% Tween 20).

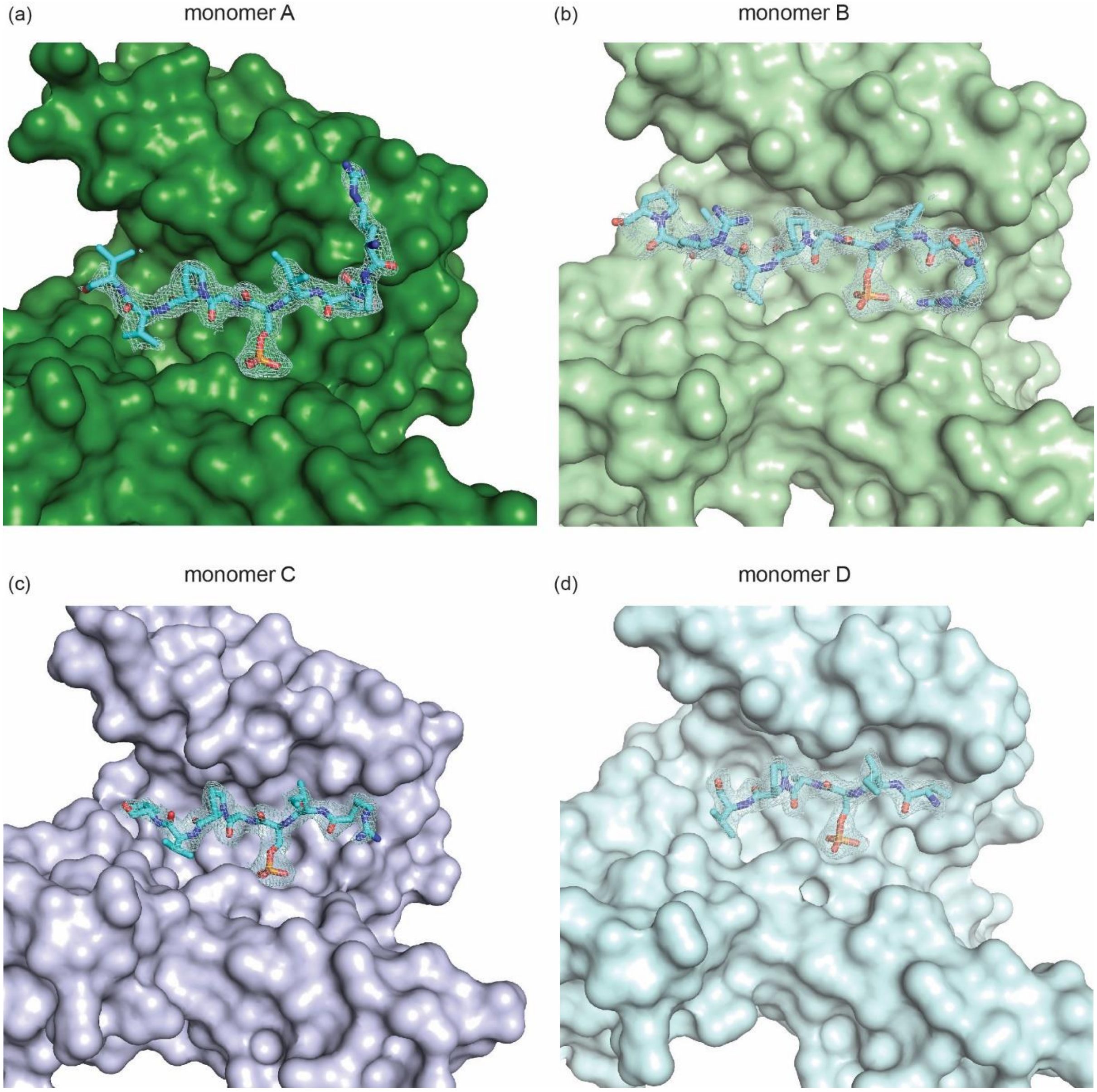

**Figure S11** Additional images for the *h*DMX$_{361\text{-}374}^{pSer367}$/14-3-3σ structure (PDB: 6YR5); each panel shows each singly phosphorylated peptide ($2F_o$- $F_c$ electron density map, contoured at 1σ) bound to a 14-3-3σ monomer (dark green, light green, light blue or light lilac surface), in its conserved amphipathic groove (*h*DMX$_{361\text{-}374}^{pSer367}$ shown as sticks, carbon in cyan, phosphorous orange, nitrogen dark blue and oxygen red)

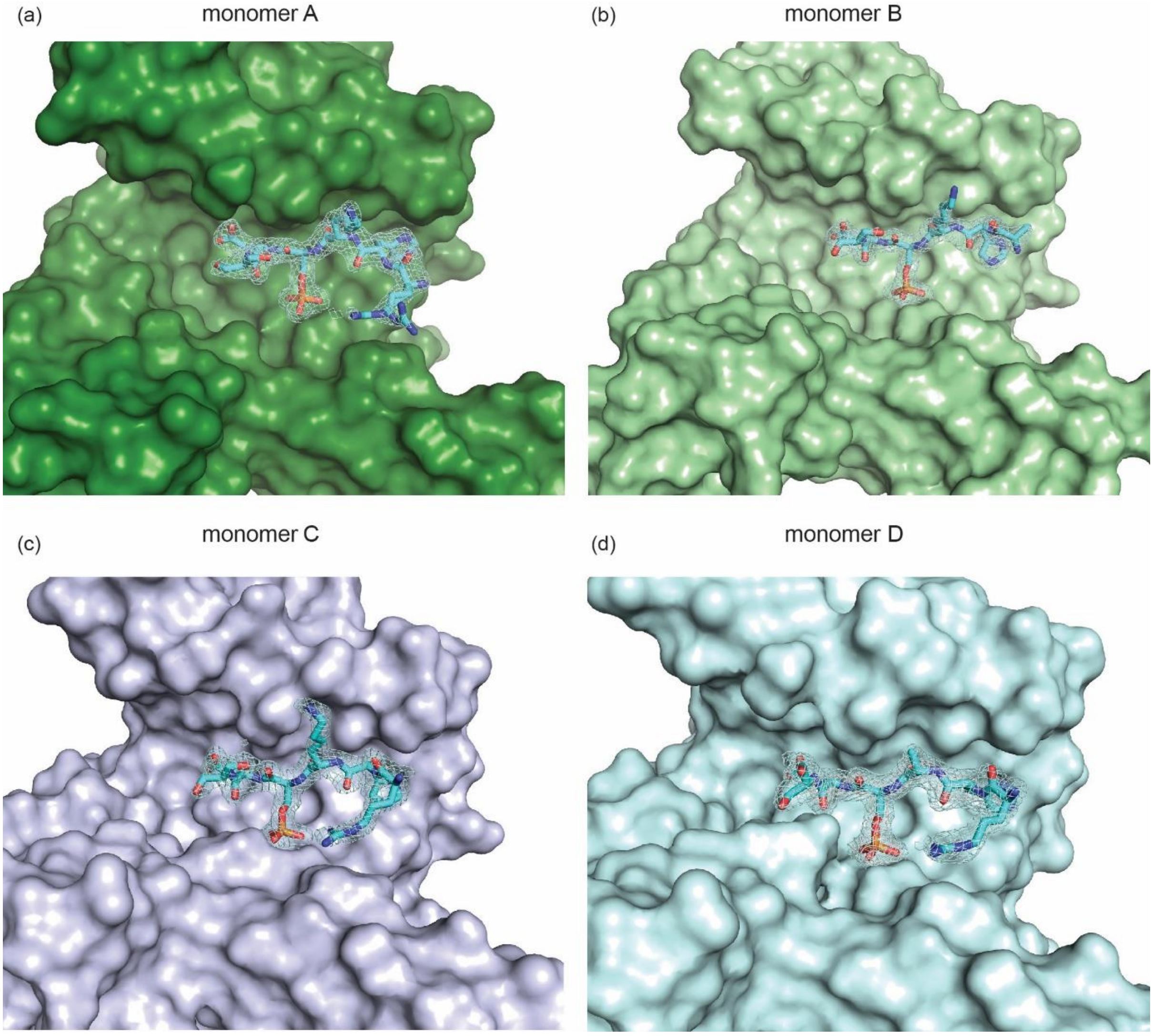

**Figure S12** Additional images for the *h*DM2$_{180\text{-}192}$$^{pSer186}$/14-3-3σ structure ((PDB: 6YR6)) each panel shows each singly phosphorylated peptide ($2F_o$- $F_c$ electron density map, contoured at 1σ) bound to a 14-3-3σ monomer (dark green, light green, light blue or light lilac surface), in its conserved amphipathic groove (*h*DM2$_{180\text{-}192}$$^{pSer186}$ shown as sticks, carbon in cyan, phosphorous orange, nitrogen dark blue and oxygen red)

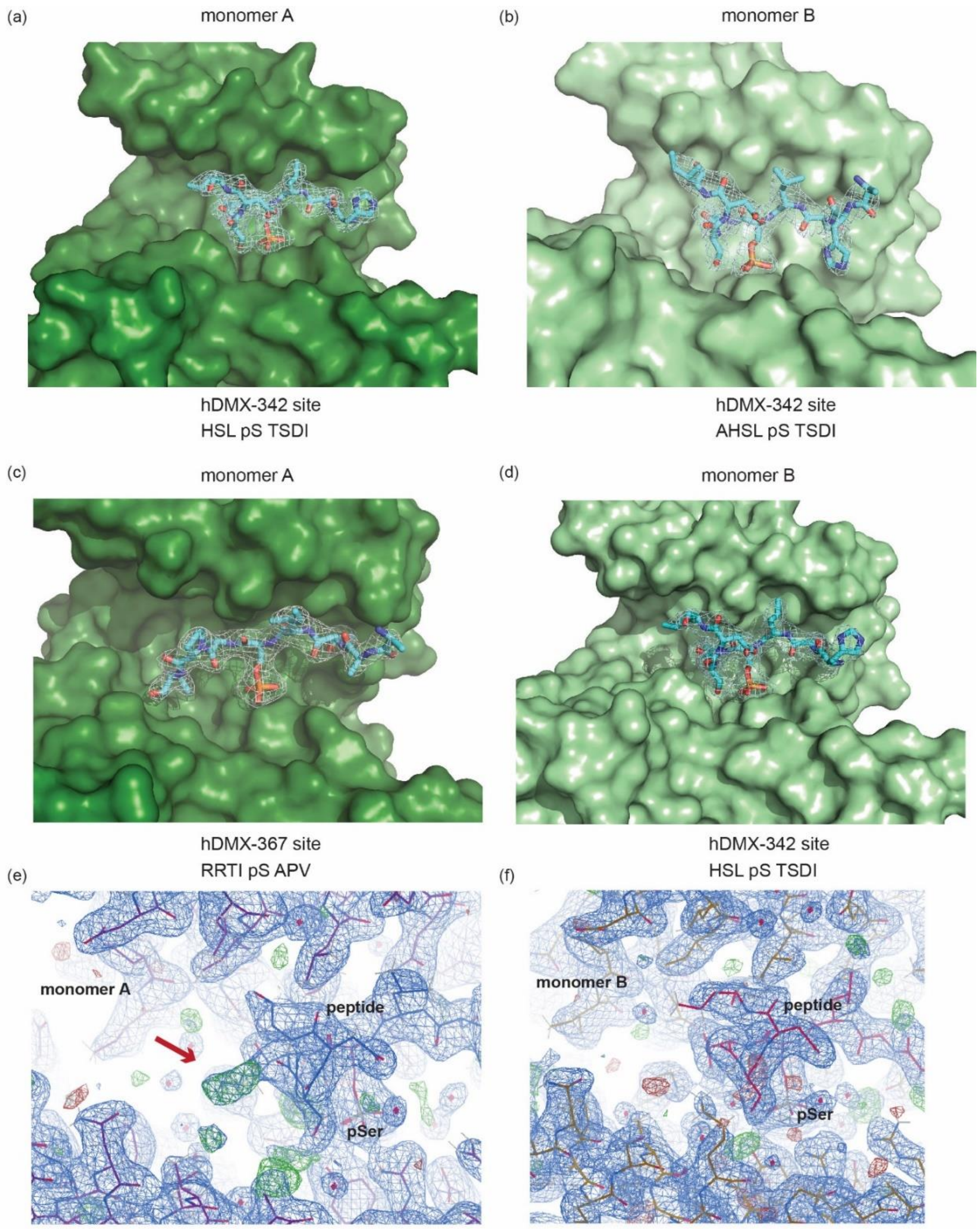

**Figure S13** *h*DMX$_{335\text{-}374}^{pSer342/pSer367}$/14-3-3σ structure (PDB ID: 6YR7) (a-b) show the final refined structure with both pSer342 in 14 3 3σ monomer A and B (dark green, light green surface), in its conserved amphipathic groove (*h*DMX-342 and *h*DMX-367 sites shown as sticks with carbon in cyan, phosphorous orange, nitrogen dark blue and oxygen red), (c-d) shows electron density during the refinement, where pSer367 could be in monomer A. (e-f) show electron density for the final refined structure ($2F_o$- $F_c$ electron density map in blue, contoured at 1σ). Extra electron density can be observed in monomer A ($F_o$- $F_c$ , contoured at 2.5σ, in green) indicating pSer342 and pSer367 peptides overlapping. For monomer A, the dominant electron density map can be assigned to the pSer342 binding site as the unusually bent conformation is not supported by the Pro369 of the second binding site (pSer367). Nevertheless, after modeling in the pSer342 site extra electron density could be observed adjacent to Ser344 of the first binding site of the peptide. This extra electron density fits to the sequence of the pSer342 binding site, confirming the compatibility of the pSer342 with monomer A.

**Table S1**. Data collection refinement statistics

| | *h*DMX$_{361-374}$$^{pSer367}$ | *h*DMX$_{345-374}$$^{pSer342/pSer367}$ | *h*DM2$_{180-192}$$^{pSer186}$ |
|---|---|---|---|
| PDB ID | 6YR5 | 6YR6 | 6YR7 |
| Space group | *P* 1 | C 1 2 1 | *P* 1 |
| Cell constants a,b,c, α, β, γ | 63.95Å 75.30Å 78.39Å 95.15◦ 113.14◦ 93.84◦ | 133.11Å 70.21Å 80.63Å 90.00◦ 101.97◦ 90.00◦ | 63.23Å 74.57Å 77.97Å 98.43◦ 111.09◦ 93.12◦ |
| Resolution (Å) | 74.53 - 2.10<br>71.50 - 2.25 | 56.26 - 2.10<br>65.11 - 2.10 | 56.90 - 1.75<br>73.26 - 1.75 |
| % Data completeness (in resolution range) | 97.3 (74.53-2.10)<br>97.6 (71.50-2.25) | 99.1 (56.26-2.10)<br>91.2 (65.11-2.10) | 94.9 (56.90-1.75)<br>90.0 (73.26-1.75) |
| $R_{merge}$ | 0.11 | 0.04 | 0.07 |
| < I/σ(I) > | 5.36 (at 2.25Å) | 0.68 (at 2.10Å) | 1.33 (at 1.75Å) |
| Refinement program | REFMAC 5.8.0230 | PHENIX 1.12_2829 | PHENIX 1.12_2829, REFMAC 5.8.0238 |
| R, $R_{free}$ | 0.198, 0.222<br>0.206, 0.230 | 0.212, 0.246<br>0.213, 0.246 | 0.199, 0.224<br>0.198, 0.223 |
| $R_{free}$ test set | 2993 reflections (4.85%) | 2108 reflections (5.02%) | 6318 reflections (5.03%) |
| Wilson B-factor (Å) | 27.3 | 35.8 | 25.3 |
| Anisotropy | 0.338 | 0.397 | 0.241 |
| Bulk solvent $k_{sol}$(e/Å$^3$), Bsol(Å$^2$) | 0.30, 31.7 | 0.36, 40.0 | 0.41, 47.8 |
| L-test for twinning | < \|L\| > = 0.47, < L2 > = 0.30 | < \|L\| > = 0.50, < L2 > = 0.34 | < \|L\| > = 0.50, < L2 > = 0.33 |
| Estimated twinning fraction | No twinning to report. | No twinning to report. | No twinning to report. |
| Fo,Fc correlation | 0.93 | 0.94 | 0.94 |
| Total number of atoms | 7709 | 7376 | 15129 |
| Average B, all atoms (Å$^2$) | 34.0 | 54.0 | 40.0 |

### Experimental Methods

#### Peptide synthesis

General remarks: Resins and amino acids were purchased from either Sigma–Aldrich or Novabiochem. All amino acids were N -Fmoc protected and side chains protected with Boc (His, Lys), t-Bu (Asp, Glu, Ser, Thr), Pbf (Arg), Trt (Asn, Gln). Peptides were synthesized either manually or using a microwave assisted automated peptide synthesizer (CEM Liberty Blue) on a 0.05 or 0.1 mmol scale. DMF used in peptide synthesis was of ACS grade and from Sigma–Aldrich.

Manual peptide synthesis: Manual peptide synthesis followed this cycle: swelling of a resin (20 min) in cartridge used for solid-phase synthesis, washing (DMF, 3 x 2 ml x 2 min), deprotection (Method A), and coupling of a desired amino acid (Method B), where successful coupling and deprotection were determined by a colour test (Method C). Acetylation (Method D) or coupling of a fluorescent dye (Method E) were performed prior to cleavage (Method F).

*Method A*: Deprotection N-terminal Fmoc-protecting groups were removed by adding 20% piperidine in DMF (5 × 2 mL × 2 min) and washed with DMF (5 × 2 mL × 2 min) after.

*Method B*: Manual coupling of amino acid and Ahx

The desired amino acid or Ahx (5 equiv.), DIPEA (10 equiv.) and HCTU (5 equiv.) were dissolved in DMF (2 mL) and added to the resin, followed by agitation for 1 h. Reagents were removed by filtration and the resin was washed with DMF (3 × 2 mL × 2 min).

*Method C*: Kaiser test

Successful coupling or deprotection for any residue coupled manually was determined by Kaiser test. A few resin beads were transferred into a vial and mixed with 2 drops of each of the solutions:

1) Ninhydrin (5% w/v) in ethanol

2) Phenol (80% w/v) in ethanol

3) 1 mM KCN (aq.) in pyridine (2% v/v)

The solution was heated at 100 °C for five minutes before observing the change in colour. Successful deprotection was observed by colour of the beads changing into blue, where successful coupling gave no change in colour.

*Method D*: N-terminal acetylation

Acetic anhydride (10 equiv.) and DIPEA (10 equiv.) were dissolved in DMF (2 mL) and the solution was transferred to the resin. After 2 h, the resin was drained and washed with DMF (3 × 2 mL × 2 min). Successful capping was determined by colour test (Method C).

*Method E*: N-terminal Fluorescent Dye coupling

5,6-carboxyfluorescein (5 equiv.), DIPEA (5 equiv.) and HCTU (5 equiv.) were dissolved in DMF (2 mL) and added to the resin, followed by agitation for 1 h. Reagents were filtered and the resin was washed with DMF (3 × 2 mL × 2 min) ahead of cleavage and deprotection.

*Method F*: Cleavage and deprotection of peptides of the resin

After elongation and acetylation or fluorescent dye coupling was complete, the resin was washed with $CH_2Cl_2$ (5 × 2 mL × 2 min), $Et_2O$ (5 × 2 mL × 2 min) and dried under vacuum. Peptides were cleaved and side-chain deprotected using 'Reagent K' TFA:EDT:Thioanisole:Phenol:$H_2O$ 82:3:5:5:5 (2 mL × 3 h). The peptide was precipitated in ice-cold $Et_2O$ (10 mL) and placed in a centrifuge (3000 rpm × 5 min). The supernatant was removed, the precipitate resuspended in ice-cold $Et_2O$ and placed in a centrifuge again (3x). The precipitate was dried under a stream of nitrogen overnight, before being dissolved in $H_2O$ and lyophilized.

<u>Automated peptide synthesis method:</u> Peptides prepared using automated peptide synthesizer followed cycles described below. Resin loading cycle cleans the reaction vessel, washes with DMF:DCM (1:1), transfers resin to reaction vessel, washes with DMF:DCM (1:1), and drains the vessel at the end of a cycle. Deprotection and coupling cycle consist of: washing with DMF (4 ml), adding 20% piperidine in DMF (6 ml), microwave deprotection cycle (30 sec), washing with DMF (4+4+4+4 mL), addition of amino acid (2.5 ml, 5 eq or 3 eq for phosphorylated amino acids), coupling reagent (1 ml, 5 eq) and activator base (0.5 ml, 5 eq), coupling microwave cycle (5 min), washing with DMF (2 ml) and draining. HCTU and DIPEA were used during automated peptide synthesis as well. As a rule, all amino acids were coupled using 75°C coupling and deprotection cycles up to Ser(PO(OBzl)OH)-OH or Thr(PO(OBzl)OH)-OH, where conventional coupling and deprotection method was used (coupling at the rt for 2 h, deprotection at rt for 15 min) as well for every amino acid following pSer/pThr. After the final residue was coupled, the resin was ejected from the reaction vessel. Ahx coupling, deprotection, acetylation or florescent dye coupling, and cleavage was performed manually using methods described above.

<u>Peptide purification:</u> Crude peptides were dissolved in $H_2O$ or DMSO and purified by UV- or MS- directed HPLC. Jupiter Proteo (250 x 21.2 mm) or a Kinetex EVO C18 (250 x 21.2 mm)

preparative column (reversed phase) was used with increasing gradient of acetonitrile in water with 0.1% formic acid, over 30 min at the flowrate of 10 ml/min. Fractions containing peptide were combined, concentrated, and lyophilized. Purity of peptides was assessed by analytical HPLC and HRMS.

#### Protein expression and purification

The pProEx HTb-His-14-3-3 constructs were expressed in BL21(DE3) cells. A single colony from a freshly made agar plate (8 g LB broth mixed with 8 g agar in 400 ml, autoclaved and poured into petri dishes to use for transformation of plasmids) was picked and mixed with 5 ml of LB media with ampicillin to inoculate a starter culture overnight at 37°C. The cells were grown in 2 L of TB media (48 g peptone, 24 g yeast, 4.6 g $KH_2PO_4$, 24 g $KHPO_4$, 5 ml glycerol in 2 L of $dH_2O$, autoclaved for 20 min at 121°C) at 37°C until the OD reached 0.4-0.6. Expression was induced by adding 0.4 mM IPTG and agitating overnight at 18°C. The expression culture was spun down (8000 rpm, 20 min, 4°C), resuspended in 200 ml of lysis buffer (50 mM Tris, 300 mM NaCl, 12.5 mM imidazole, 2 mM β-mercaptoethanol) with 5 mM $MgCl_2$ and DNAse (1:1000). The cells were lysed by French press or sonication and the solid fragments were removed by centrifugation (20000 rpm, 30 min, 4°C). The cleared lysate was loaded on a $Ni^{2+}$-NTA column, washed with 50 mM Tris, 300 mM NaCl, 12.5 mM imidazole, 2 mM β-mercaptoethanol, 0.1% triton X-100, and the protein was eluted with 50 mM Tris, 300 mM NaCl, 250 mM imidazole, 2 mM β-mercaptoethanol. Imidazole was removed by overnight dialysis using the Tris buffer (50 mM Tris, 300 mM NaCl, 2 mM β-mercaptoethanol), full length proteins were concentrated by centrifugation, rebuffered in HEPES buffer (25 mM HEPES pH 7.5, 100 mM NaCl, 10 mM $MgCl_2$, 0.5 mM TCEF) and stored in -80°C freezer. ΔC proteins were dialyzed with TEV protease to remove the expression tag and purified again on $Ni^{2+}$-NTA column, followed by size-exclusion chromatography (HiLoad 16/600 Superdex 75 pg column) in HEPES buffer (20 mM HEPES, 150 mM NaCl, 2 mM DTT). ΔC proteins were concentrated and rebuffered for -80°C storage. All proteins were analyzed by ESI QTOF-MS.

#### Fluorescence anisotropy

Direct titration assay: All assays were performed in 384 well plates (each experiment was run in triplicates) and data were collected by Perkin Elmer EnVision 2013 plate reader with excitation at 480 nm (30 nm bandwidth), polarised dichroic mirror at 505 nm and emission at 535 nm (40 nm bandwidth, S and P polarised). Experiments were carried out in HBS buffer (10 mM HEPES, 150 mM NaCl, pH 7.4) + 0.1% Tween 20 + 0.1% BSA). A ½ fold dilution series was performed in the titration, with plates read after 30 min, 4 h and 20-24 h. These gave consistent data and values when fitted. Collected data were processed in Microsoft Excel using the equations bellow. Total intensity I and anisotropy r were calculated using Equations 1 and 2 for each well. Average anisotropy was plotted against protein concentration using

OriginPro and logistic curve was fitted to give $r_{min}$ and $r_{max}$. Using Equation 3 anisotropy was converted into fraction bound and multiplied by peptide concentration to be fitted in Origin using Equation 5 to obtain $K_d$ values.

Equation 1. $I = 2PG + S$

Equation 2. $r = \frac{S-PG}{I}$

Equation 3. $L_b = \frac{(r-r_{min})}{\lambda(r_{max}-r)+r-r_{min}}$

Equation 4. $y = r_{min} + \frac{r_{max}-r_{min}}{1+10^{(x-logx_0)}}$

Equation 5. $y = \frac{((K+X+FL)-\sqrt{((K+X+FL)^2-4xFL))}}{2}$

r = anisotropy, I = total intensity, P = perpendicular intensity, S = parallel intensity, $L_b$ = fraction ligand bound, $\lambda = I_{bound}/I_{unbound} = 1$, FL = fluorescent ligand concentration, K = $K_d$

#### Surface plasmon resonance

Experiments were performed at 25°C using a Biacore T200 instrument (GE Healthcare). For immobilization, the running buffer was HBS buffer (HEPES, NaCl, Tween-20, pH 7.4). 14-3-3η was immobilized on an NTA sensor chip (Series S Sensor Chip GE Healthcare) using NTA reagent and Biacore Amine Coupling Kit. Briefly, 14-3-3η was diluted to 0.2 mg/ml and 200 µL was injected over a chip surface that had been activated with an injection of $Ni^{2+}$, followed by 140 µL of 1:1 NHS/EDC. 30 µL of 0.5 M ethanolamine was then injected to cap the excess free amine groups. Immobilization levels of the 14-3-3η were found to be around 2000 response units (RU) for kinetic measurements. *h*DMX and *h*DM2 peptides were serially diluted 11 times from the concentrations 10 times the $K_d$ values measured in FA, on a 96-well plate. Then, 20 µL of these solutions were injected over a 14-3-3η immobilized surface for 1 min at the 20 ml/min flowrate, followed by a 4 min regeneration period with HBS buffer. Kinetics and the binding affinity were calculated by using the Biacore evaluation software. Standard dose response curves were fitted by plotting RUs against the fragment concentration.

#### Isothermal titration calorimetry

ITC experiments were carried out on a MicroCal iTC200 in HBS buffer at 25℃ (25 mM HEPES pH 7.5, 100 mM NaCl, 10 mM $MgCl_2$, 0.5 mM TCEP). Peptides (0.2-1 M) in the syringe were titrated into a cell containing 14-3-3η (0.02 or 0.1 M). Data was fitted by using MicroCal software to give binding constant ($K_d$), enthalpy (ΔH), entropy (ΔS) and binding stoichiometry (n).

#### Protein crystallography

A solution of 14 mg/ml 14-3-3σΔC with *h*$DMX_{361-374}^{pSer367}$ peptide in 1:2 molar ratio was incubated overnight in crystallization buffer (20 mM HEPES pH 7.5, 2 mM $MgCl_2$, and 2 mM

BME). Crystals were obtained at 4°C in a hanging drop vapour diffusion set up with 0.2 M sodium sulfate, 0.1 M Bis-Tris propane, 20 % w/v PEG 3350 at pH 8.5 as precipitation buffer. Protein/peptide solution and precipitation buffer were mixed in a 1:1 ratio in a total volume of 2 µL. Diffraction data was recorded at the DESY, Petra III P11, Hamburg to a resolution of 2.2 Å. $h\text{DMX}_{335\text{-}374}^{\text{pSer342/pSer367}}$ crystals were obtained at 4°C after overnight incubation of 12 mg/ml 14-3-3σ ΔC with $h\text{DMX}_{335\text{-}374}^{\text{pSer342/pSer367}}$ peptide (1:2 ratio) in sodium citrate dihydrate, bis-tris propane, pH 8, 20% PEG 3350. $h\text{DM2}_{180\text{-}192}^{\text{pSer186}}$ crystals were obtained at 4°C after overnight incubation of 12 mg/ml 14-3-3σ ΔC with $h\text{DM2}_{180\text{-}192}^{\text{pSer186}}$ (1:2 ratio) in 0.0375 M $CdSO_4$ $H_20$, 0.075 M HEPES, pH 7.5, 0.75 M NaAc $3H_20$, 25% glycerol. Diffraction data for $h\text{DMX}_{335\text{-}374}^{\text{pSer342/pSer367}}$ and $h\text{DM2}_{180\text{-}192}^{\text{pSer186}}$ crystals were recorded at the Diamond Light source, UK with a resolution of 2.1 Å for $h\text{DMX}_{335\text{-}374}^{\text{pSer342/pSer367}}$ and a resolution of 1.75 Å for $h\text{DM2}_{180\text{-}192}^{\text{pSer186}}$. The CCP4 software package[1] was used for structure determination by molecular replacement with 4DAT (PDB accession code)as a template for the 14-3-3σ structure. Further rounds of manual model building and refinement were performed using COOT[2] and REFMAC[3] respectively. Data collection and refinement statistics for each structure are provided in Supplementary info (Table A.4).

#### Analytical ultracentrifugation

AUC-SV experiments were performed using a Beckman Coulter Optima XL-I ultracentrifuge. Samples were prepared in HBS buffer (10 mM HEPES, 150 mM NaCl, pH 7.4) and loaded into 12 mm aluminium centrepieces with sapphire windows. Three samples were run: I. 14 µM 14-3-3η, II. 14 µM 14-3-3η + 7 µM $h\text{DMX}_{335\text{-}374}^{\text{pSer342/pSer367}}$, III. 14 µM 14-3-3η + 14 µM $h\text{DMX}_{335\text{-}374}^{\text{pSer342/pSer367}}$, with buffer as a reference, and recorded at 48 000 rpm in the An50-Ti rotor, at the 25°C. The diffusion-deconvoluted sedimentation coefficient distributions c(s) were calculated using the SEDFIT program.

#### Circular dichroism (CD) spectroscopy

Spectra were recorded on a Chirascan circular dichroism spectropolarimeter (Applied Photophysics), using 1 mm cells, scan speed of 5 nm/min, 2 nm bandwith, and 180 nM to 260 nM range. The experiments were preformed in 50 mM sodium phosphate buffer, pH 7.5 at 20°C. The spectra were averaged over three repeats with a buffer baseline subtracted. Protein concentrations of approximately 0.2 mg/mL were used for all proteins.

### Peptide and protein analytical characterization

**Table S2**. High resolution mass spectrometry data for peptides

| Peptide | $[M+2H]^{1+}$ Obs[d] | $[M+2H]^{1+}$ Exp[d] | $[M+3H]^{2+}$ Obs[d] | $[M+3H]^{2+}$ Exp[d] | $[M+4H]^{3+}$ Obs[d] | $[M+4H]^{3+}$ Exp[d] |
|---|---|---|---|---|---|---|
| *h*DMX$_{144\text{-}158}^{pThr151}$ | 920.9451 | 920.9437 | 614.2984 | 614.2982 | N/A | 460.9755 |
| FAM-Ahx *h*DMX$_{144\text{-}158}^{pThr151}$ | 1136.0173 | 1136.0086 | 757.6755 | 757.6748 | 568.5060 | 568.5079 |
| FAM-Ahx *h*DMX$_{144\text{-}158}$ | 1096.0239 | 1096.0255 | 731.0186 | 731.0194 | 568.5060 | 548.5163 |
| *h*DMX$_{335\text{-}349}^{pSer342}$ | 847.9230 | 847.9217 | 565.6169 | 565.6169 | N/A | 424.4645 |
| FAM-Ahx *h*DMX$_{335\text{-}349}^{pSer342}$ | 1062.9899 | 1062.9866 | 708.9934 | 708.9935 | N/A | 531.9970 |
| FAM-Ahx *h*DMX$_{335\text{-}349}$ | 1023.0011 | 1023.0035 | 682.3364 | 682.3381 | N/A | 512.0054 |
| *h*DMX$_{361\text{-}374}^{pSer367}$ | 859.9489 | 859.9478 | 573.6359 | 573.6343 | 430.4781 | 430.4776 |
| FAM-Ahx *h*DMX$_{361\text{-}374}^{pSer367}$ | 1075.0167 | 1075.0128 | 717.0097 | 717.0109 | 537.9939 | 538.0100 |
| FAM-Ahx *h*DMX$_{361\text{-}374}$ | 1035.0430 | 1035.0296 | 690.3517 | 690.3555 | 517.9810 | 518.0185 |
| *h*DM2$_{160\text{-}171}^{pSer166}$ | 784.8648 | 784.8625 | 523.5776 | 523.5774 | N/A | 392.9349 |
| FAM-Ahx *h*DM2$_{160\text{-}171}^{pSer166}$ | 999.9097 | 999.9274 | 666.9422 | 666.9540 | 500.2079 | 500.4674 |
| *h*DM2$_{180\text{-}192}^{pSer186}$ | 831.9259 | 831.9254 | 554.9526 | 554.9527 | 416.4659 | 416.4663 |
| FAM-Ahx *h*DM2$_{180\text{-}192}^{pSer186}$ | 1047.0032 | 1046.9904 | 698.3269 | 698.3293 | 523.9629 | 523.9988 |
| | **$[M+3H]^{1+}$ Obs[d]** | **$[M+3H]^{1+}$ Exp[d]** | **$[M+4H]^{2+}$ Obs[d]** | **$[M+4H]^{2+}$ Exp[d]** | **$[M+5H]^{3+}$ Obs[d]** | **$[M+5H]^{3+}$ Exp[d]** |
| *h*DMX$_{335\text{-}373}^{pSer342/pSer367}$ | 1479.7087 | 1479.046 | 1110.5397 | 1109.5366 | 888.4359 | 887.8308 |
| FAM-Ahx *h*DMX$_{335\text{-}373}^{pSer342/pSer367}$ | 1622.7495 | 1622.087 | 1217.3286 | 1216.8169 | 974.0712 | 973.655 |
| FAM-Ahx *h*DMX$_{345\text{-}373}^{pSer342}$ | 1595.7648 | 1595.431 | 1197.3266 | 1196.8254 | 957.8637 | 957.6617 |
| FAM-Ahx *h*DMX$_{345\text{-}373}^{pSer367}$ | 1595.7632 | 1595.431 | 1197.0930 | 1196.8254 | 957.8848 | 957.6617 |
| *h*DM2$_{160\text{-}192}^{pSer166/pSer186}$ | 1349.6392 | 1349.308 | 1012.7332 | 1012.2325 | 810.3888 | 809.9875 |
| FAM-Ahx *h*DM2$_{160\text{-}192}^{pSer166/pSer186}$ | 1493.0156 | 1492.348 | 1120.0278 | 1119.5129 | 896.2256 | 895.8117 |
| FAM-Ahx *h*DM2$_{160\text{-}192}$ | 1439.7003 | 1439.037 | 1079.7792 | 1079.5297 | 864.0253 | 863.8252 |

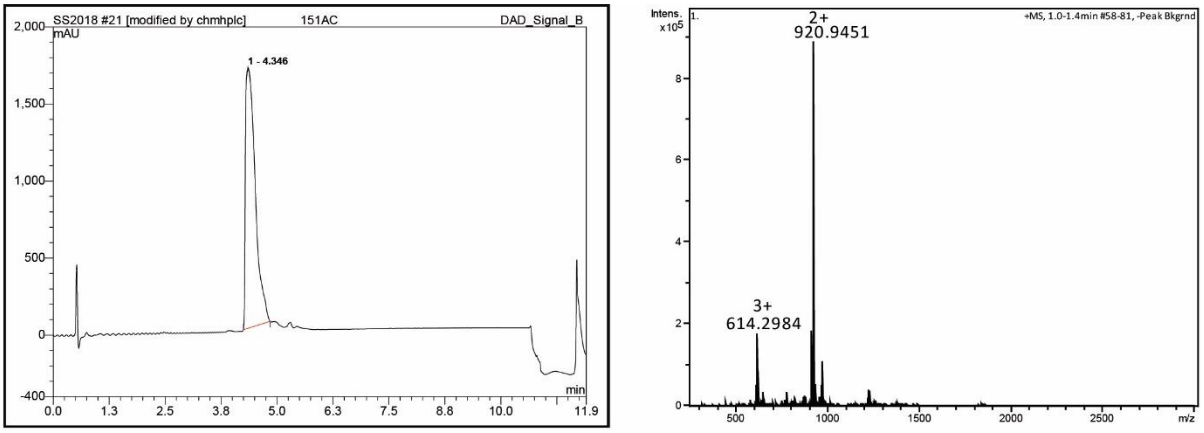

*h*DMX$_{144\text{-}158}^{pThr151}$

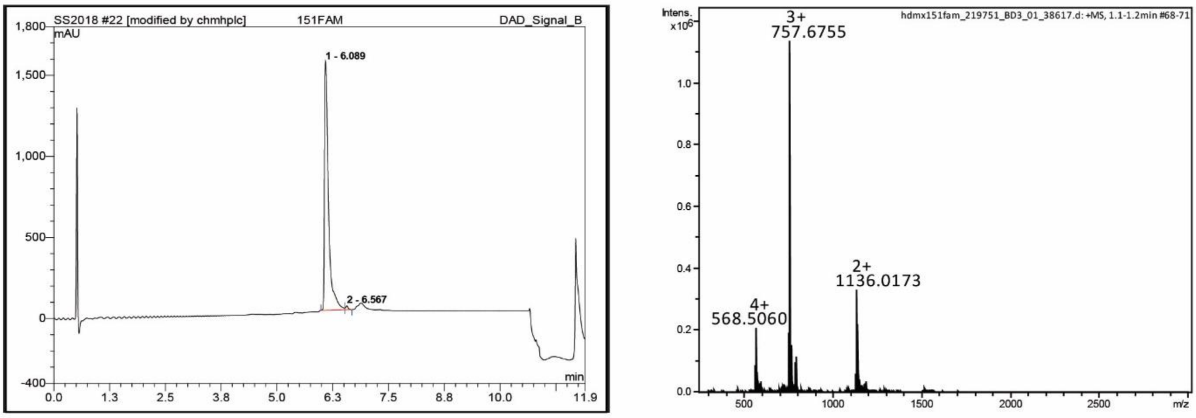

FAM-Ahx *h*DMX$_{144\text{-}158}^{pThr151}$

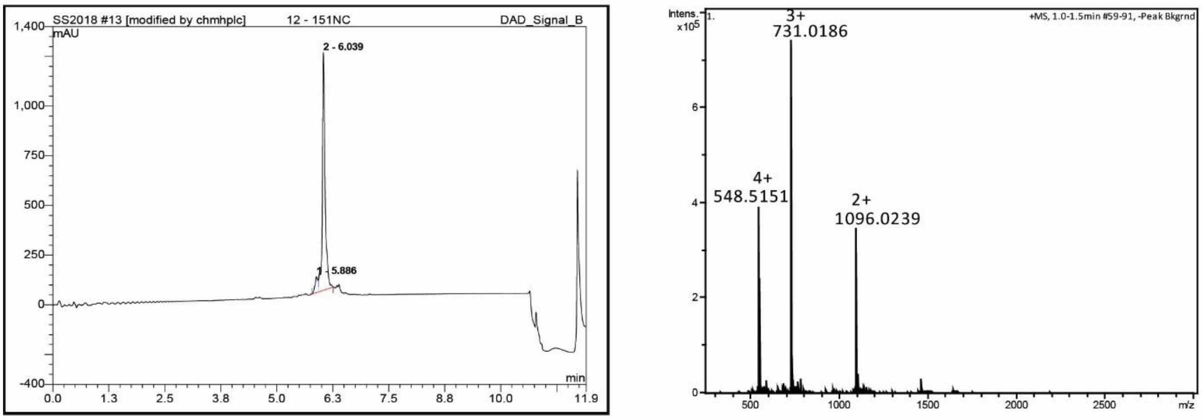

FAM-Ahx *h*DMX$_{144\text{-}158}$

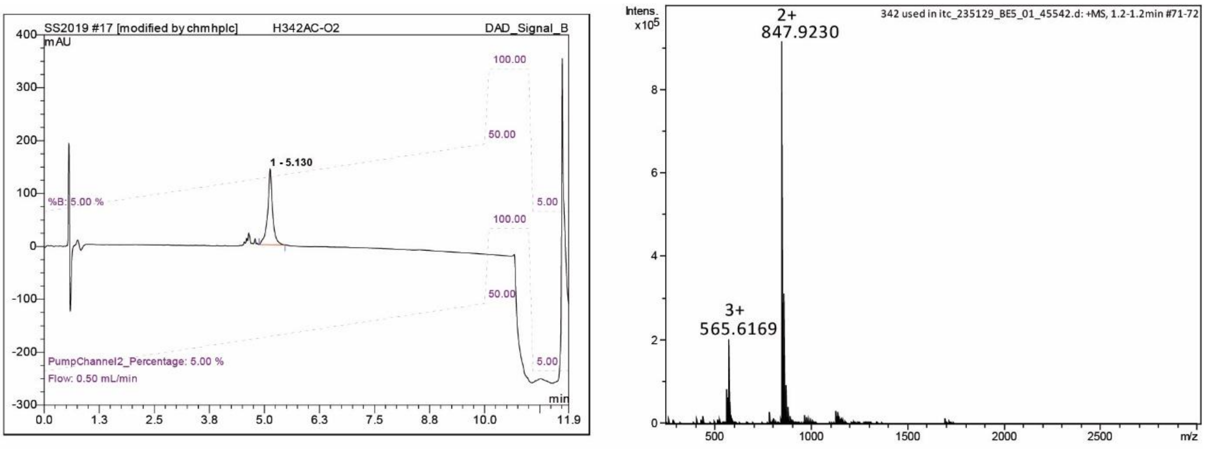

*h*DMX$_{335-349}$$^{pSer342}$

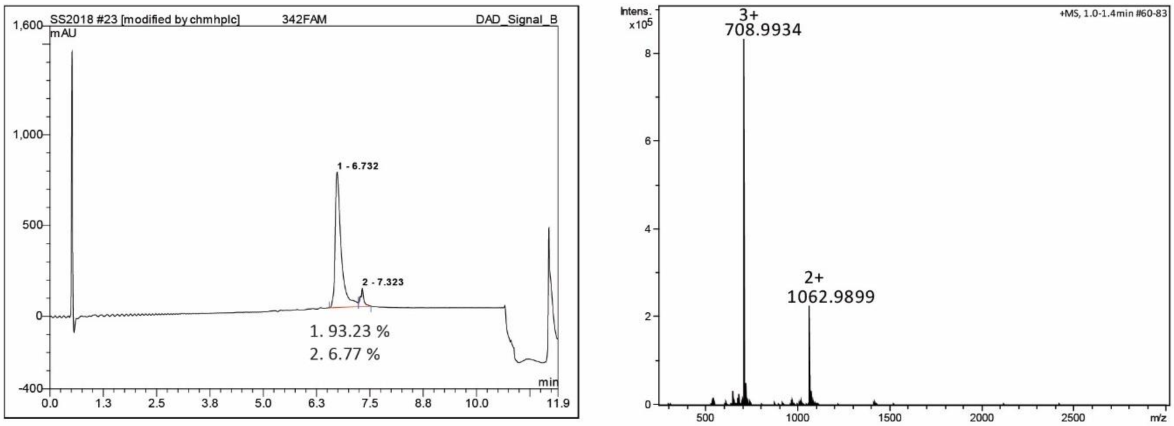

FAM-Ahx *h*DMX$_{335-349}$$^{pSer342}$

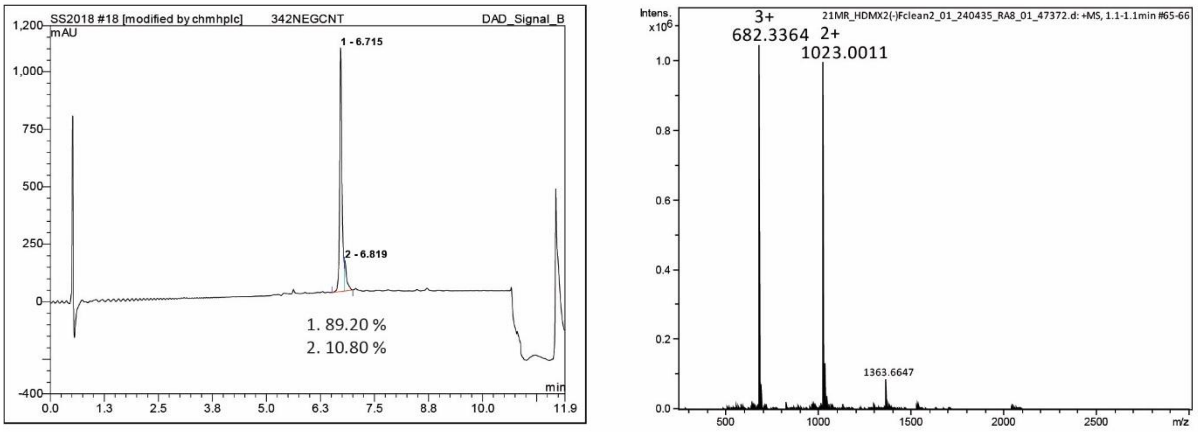

FAM-Ahx *h*DMX$_{335-349}$

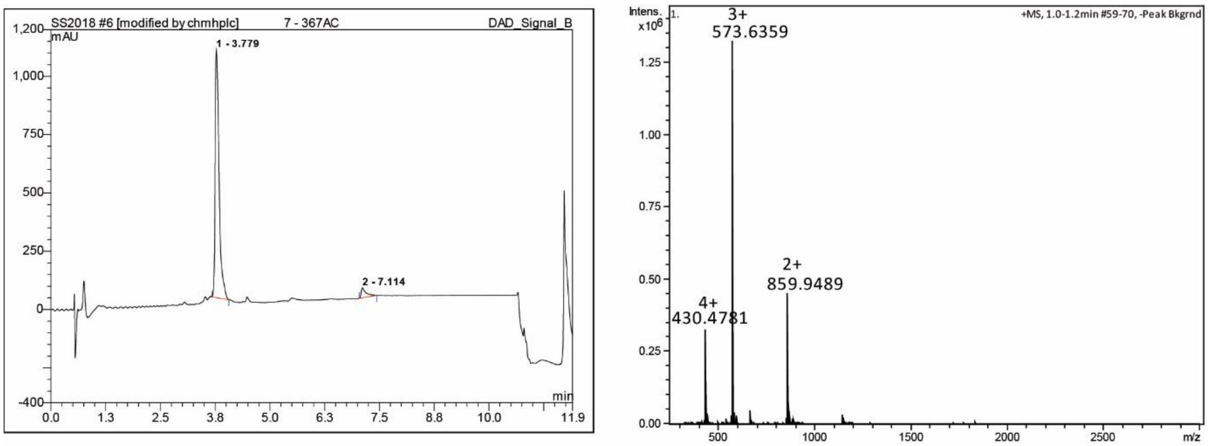

*h*DMX$_{361\text{-}374}$$^{pSer367}$

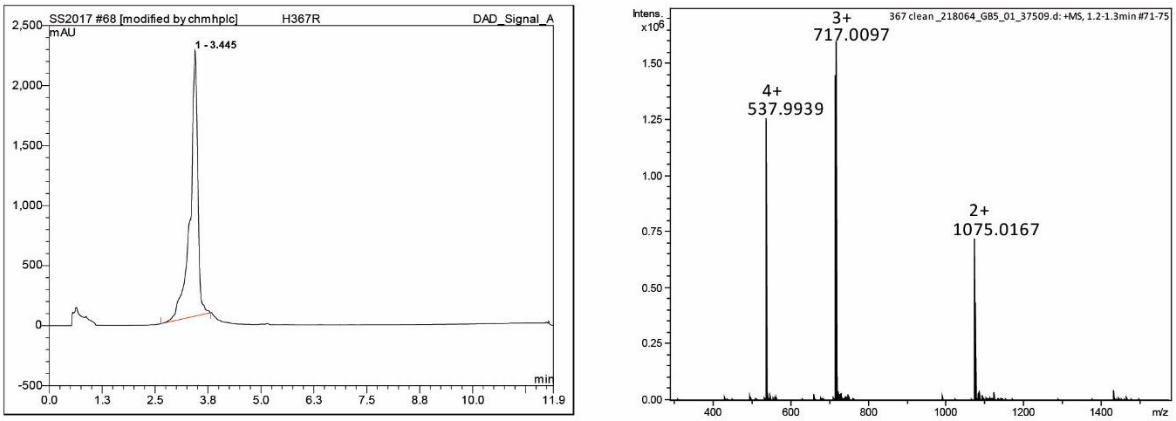

FAM-Ahx *h*DMX$_{361\text{-}374}$$^{pSer367}$

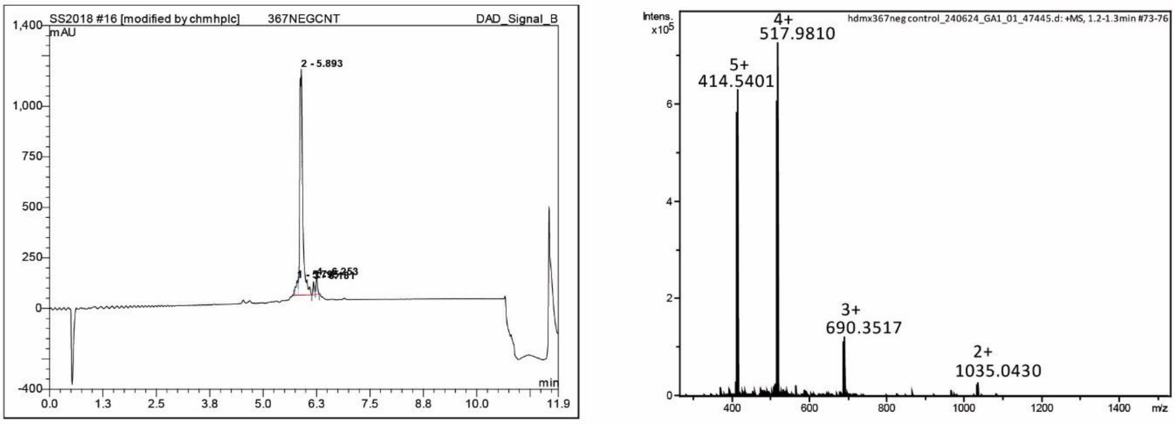

FAM-Ahx *h*DMX$_{361\text{-}374}$

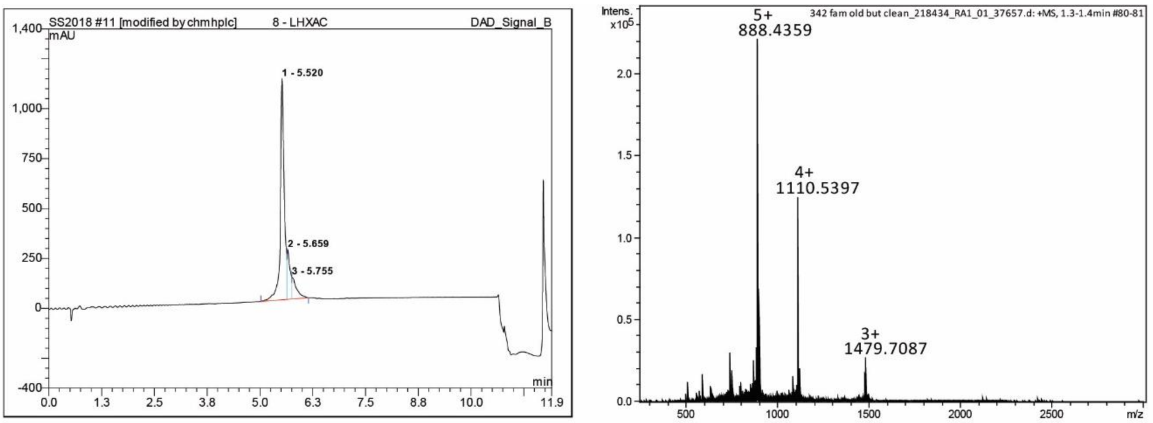

*h*DMX$_{333\text{-}373}$$^{pSer342/pSer367}$

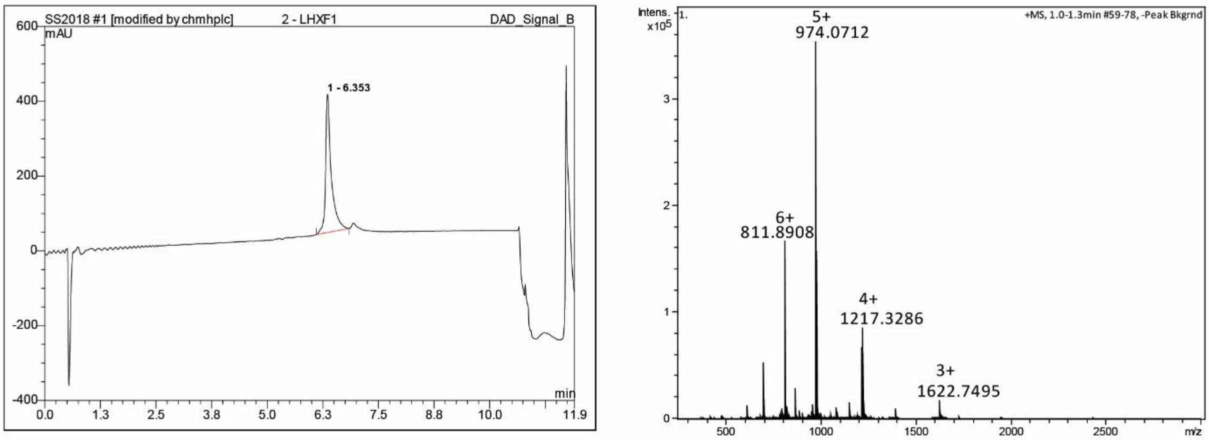

FAM-Ahx *h*DMX$_{333\text{-}373}$$^{pSer342/pSer367}$

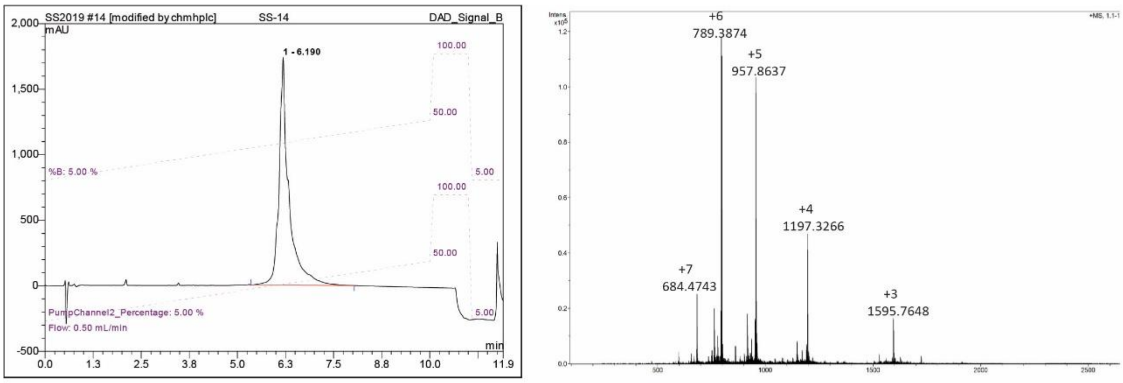

FAM-Ahx *h*DMX$_{333\text{-}373}^{pSer342}$

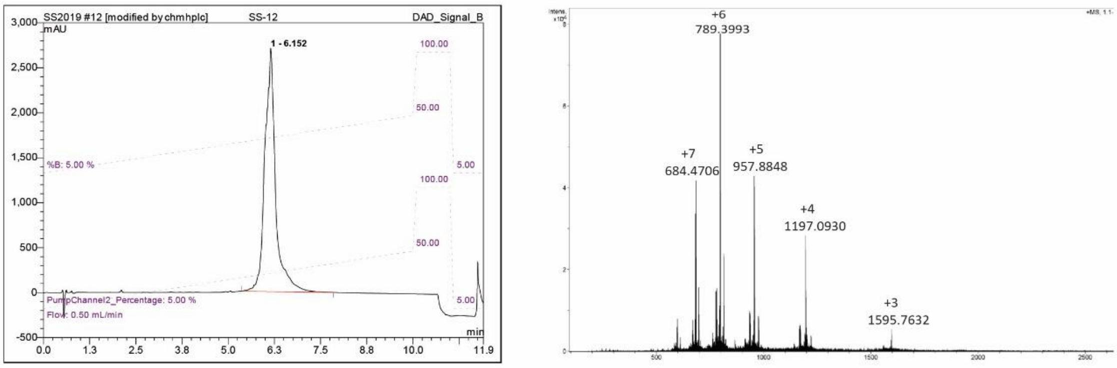

FAM-Ahx *h*DMX$_{333\text{-}373}^{pSer367}$

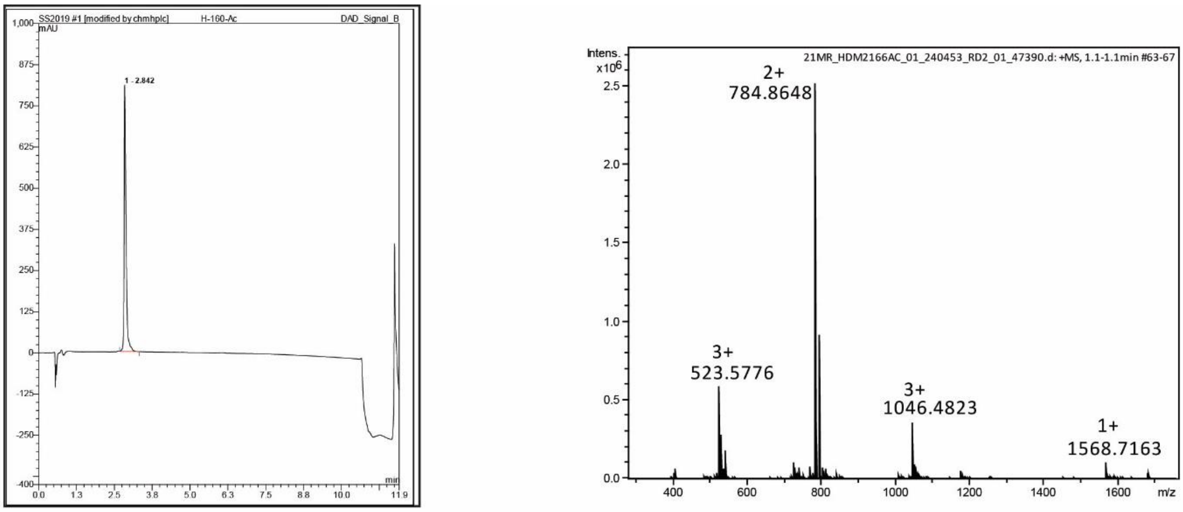

$h$DM2$_{160\text{-}171}^{\text{pSer166}}$

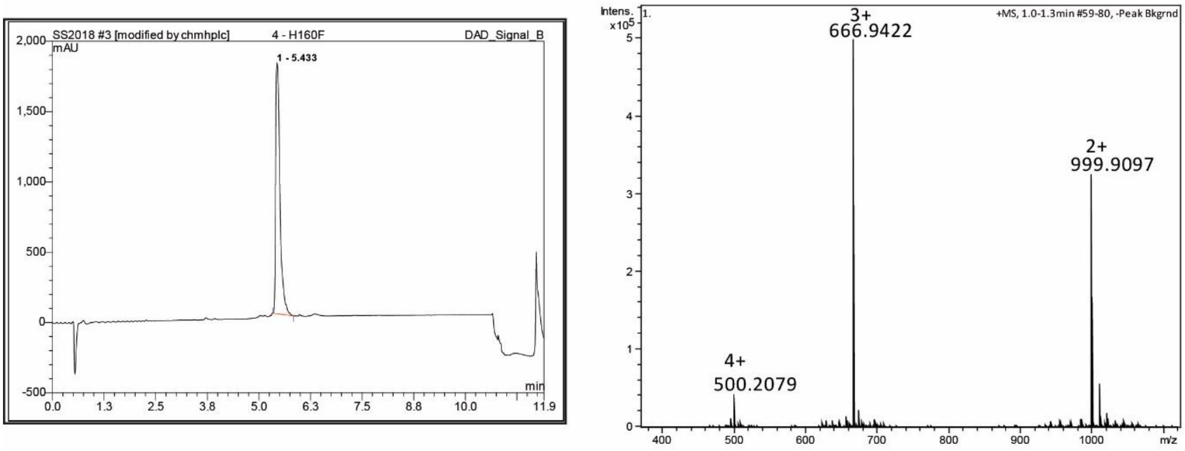

FAM-Ahx $h$DM2$_{160\text{-}171}^{\text{pSer166}}$

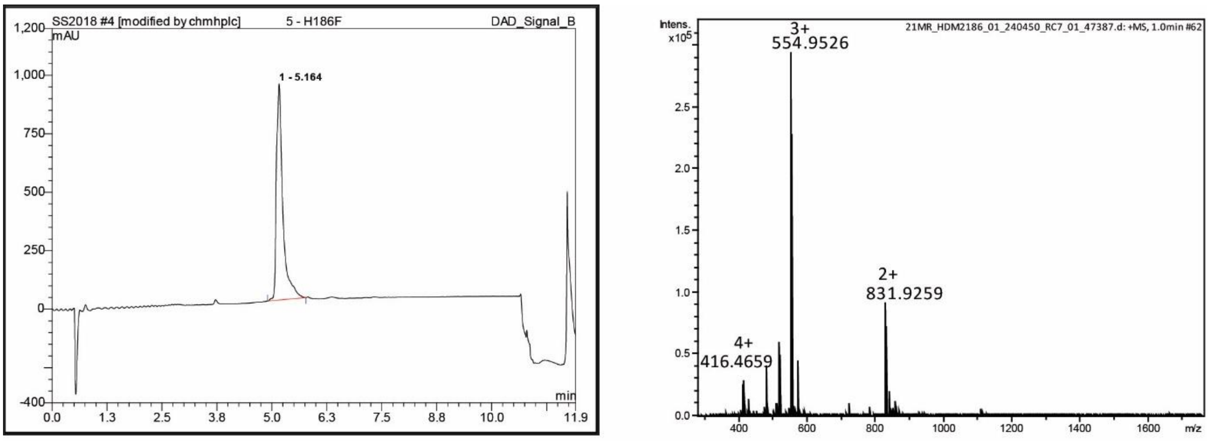

$h$DM2$_{180\text{-}192}$$^{pSer186}$

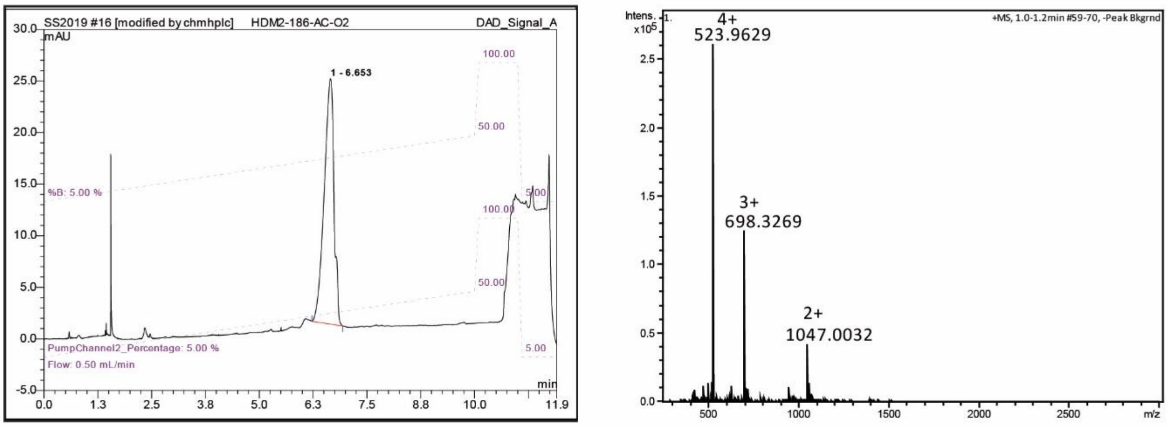

FAM-Ahx $h$DM2$_{180\text{-}192}$$^{pSer186}$

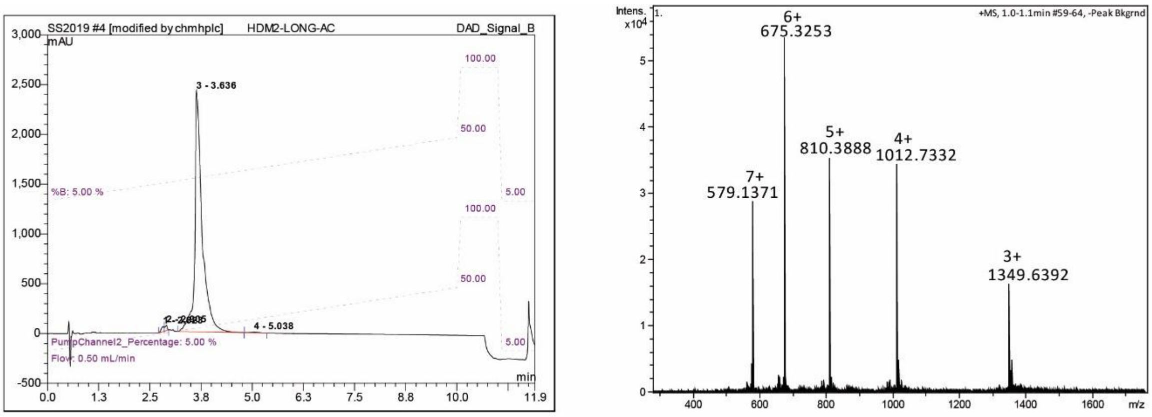

$h$DM2$_{160\text{-}192}^{\text{pSer166/pSer186}}$

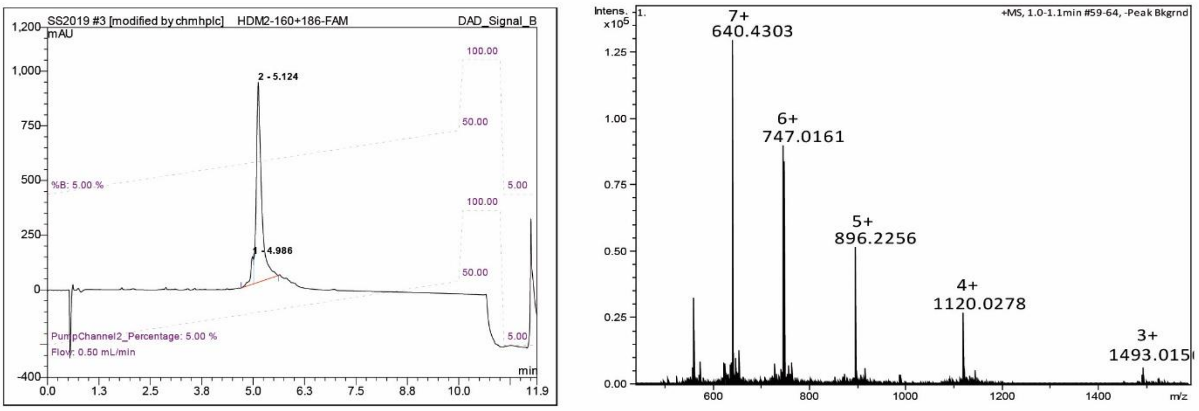

FAM-Ahx $h$DM2$_{160\text{-}192}^{\text{pSer166/pSer186}}$

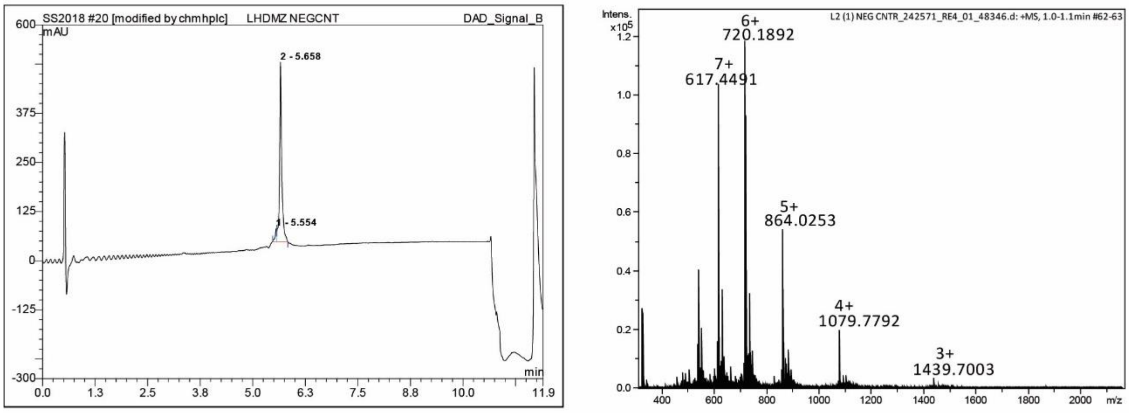

FAM-Ahx $h$DM2$_{160\text{-}192}^{\text{pSer166/pSer186}}$

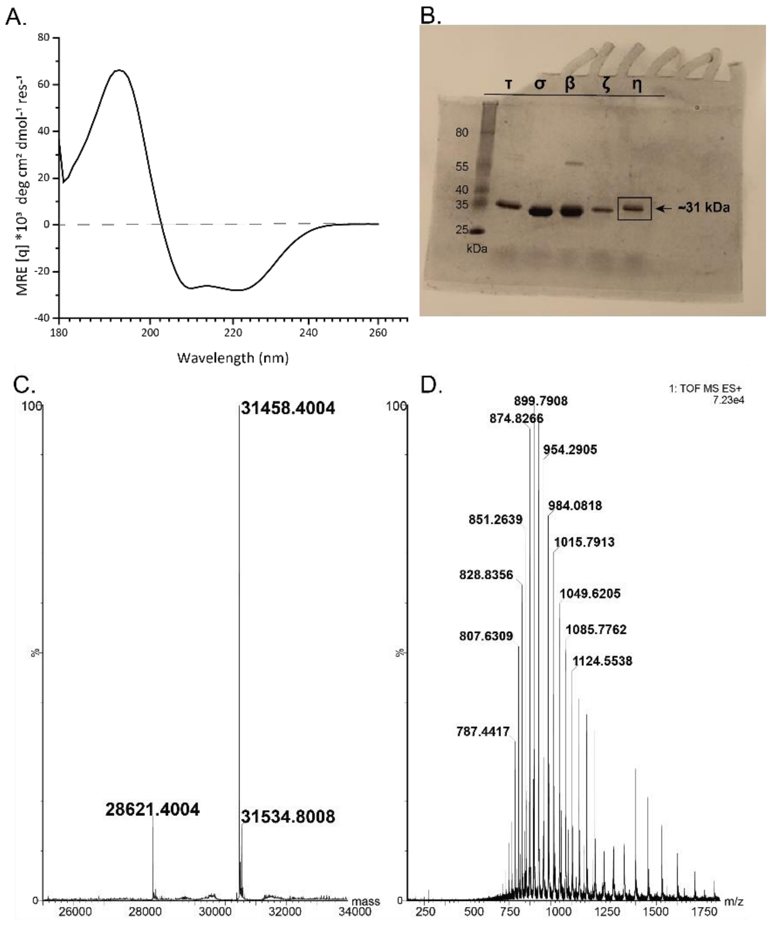

Representative 14-3-3 characterization and quality control (shown for 14-3-3η) (A) CD spectra showing an α-helical structure of the 14-3-3η (71%, theoretical helicity 78%, at 20 ℃, 50 mM sodium phosphate buffer, pH 7.5) (B) Coomassie Brilliant Blue stained SDS-PAGE gel after the protein purification C) Deconvoluted spectrum of 14-3-3η (D) QTOF MS showing a mass of 31 458.40 kDa (expected: 31 459, mass of 28 621.40 kDa corresponds to 14-3-3η without $His_6$-tag and the mass of 31 534.80 Da corresponds to addition of mercaptoethanol

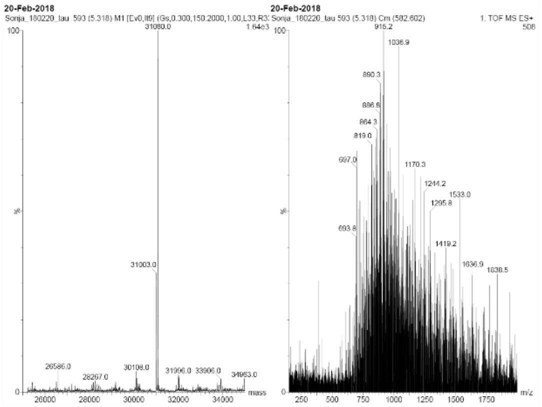

14-3-3 TAU
Mw=31005

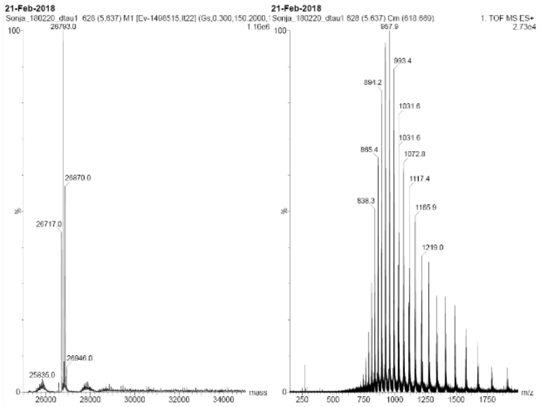

14-3-3 TAU DELTA C
Mw= 26718.3

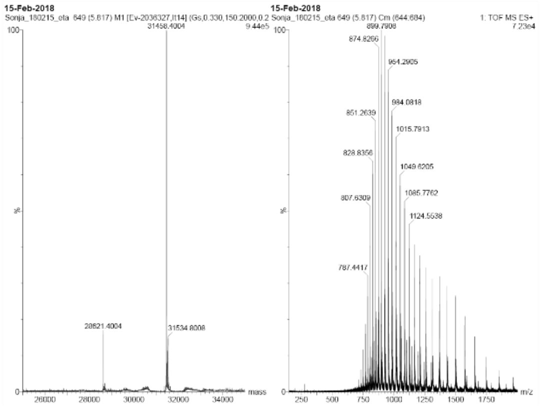

14-3-3 ETA
Mw= 31459

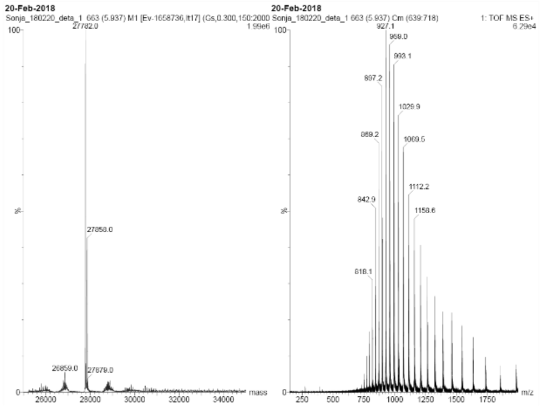

14-3-3 ETA DELTA C
Mw= 27782.4

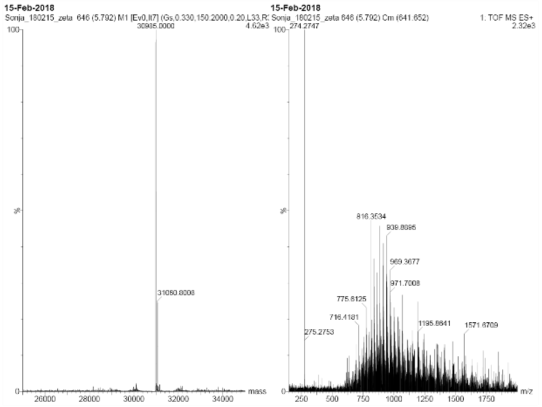

14-3-3 ZETA
Mw= 30985,5

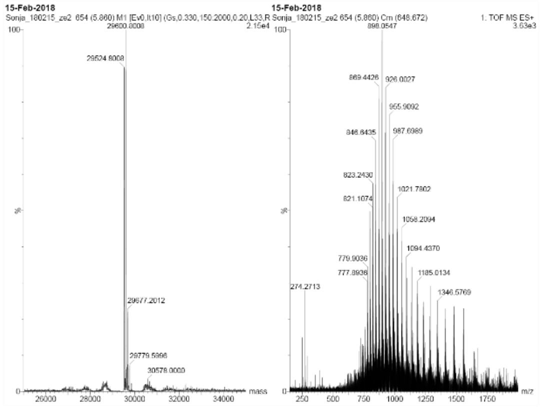

14-3-3 ZETA DELTA C
Mw= 26688.2

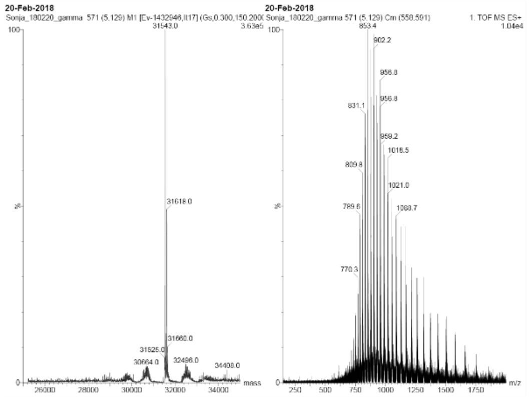

14-3-3 GAMMA
Mw= 31543

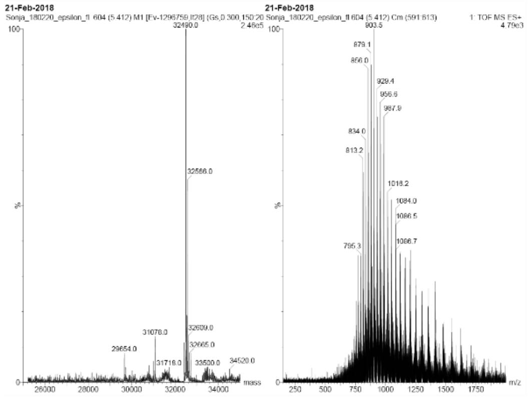

14-3-3 EPSILON
Mw= 32414

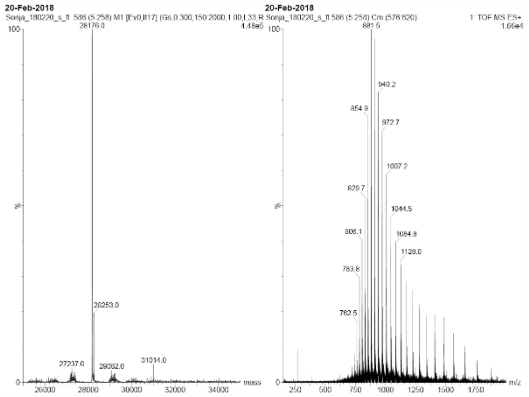

14-3-3 SIGMA
Mw= 31014.4
Major peak is 14-3-3 without his-tag

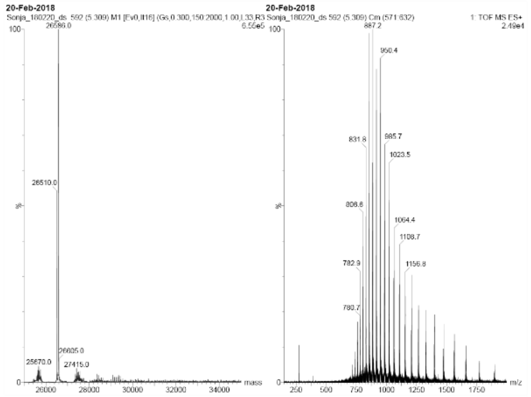

14-3-3 SIGMA DELTA C
Mw=26509.9
